# Heterogeneous identity, stiffness and growth characterise the shoot apex of *Arabidopsis* stem cell mutants

**DOI:** 10.1101/2023.07.28.550972

**Authors:** Léa Rambaud-Lavigne, Aritra Chatterjee, Simone Bovio, Virginie Battu, Quentin Lavigne, Namrata Gundiah, Arezki Boudaoud, Pradeep Das

## Abstract

Stem cell homeostasis in the shoot apical meristem involves a core regulatory feedback loop between the signalling peptide CLAVATA3, produced in stem cells, and the transcription factor WUSCHEL, expressed in the underlying organising centre. *clavata* mutants display massive meristem overgrowth, which is thought to be caused by stem cell overproliferation, although it is unknown how uncontrolled stem cell divisions lead to this altered morphology. Here we first reveal local buckling defects in mutant meristems, and use analytical models to show how mechanical properties and growth rates may contribute to the phenotype. Indeed, *clavata* meristems are mechanically more heterogeneous than the wild type, and also display regional growth heterogeneities. Furthermore, stereotypical wild-type meristem organisation is lost in mutants, in which cells simultaneously express distinct fate markers. Finally, cells in mutant meristems are auxin responsive, suggesting that they are functionally different from wild-type stem cells. Thus all benchmarks show that *clavata* meristem cells are different from wild-type stem cells, suggesting that fasciation is caused by the disruption of a more complex regulatory framework that maintains distinct genetic and functional domains at the shoot apex.

**Summary statement:** Heterogeneities in cell mechanics, growth, function and identity contribute to buckling in *clavata* mutant shoot apices.

## Introduction

The shoot apical meristem (SAM) is a structure at the growing plant tip that gives rise to all its aboveground tissues. The SAM is sustained by the activity of a small pool of centrally-located undifferentiated stem cells in the slow-dividing central zone (CZ). Through successive divisions, their daughter cells exit from the CZ into the surrounding peripheral zone (PZ), where they may differentiate into lateral organs, such as leaves or flowers. The CZ is itself maintained via the activity of the underlying organising centre (OC) (Laux et al., 1996).

The regulatory network driving stem cell maintenance in *Arabidopsis* is well studied. At its core is a feedback loop between WUSCHEL (WUS), a homeobox transcription factor expressed in the OC, and CLAVATA3 (CLV3), a signalling peptide expressed in the CZ (Brand et al., 2000; Clark et al., 1995; Laux et al., 1996; Schoof et al., 2000). WUS moves from the OC to the CZ and induces stem cell identity by directly binding to the *CLV3* promoter and activating its expression (Daum et al., 2014; Yadav et al., 2011). In turn, CLV3 binds receptors, such as the leucine-rich-repeat receptor-like kinase CLV1 (Clark et al., 1997) or the receptor-like proteins CLV2 (Kayes and Clark, 1998) and CORYNE (CRN) (Fletcher et al., 1999; Jeong et al., 1999; Miwa et al., 2008; Müller et al., 2008). Signalling from these ligand-receptor complexes ultimately leads to the downregulation of *WUS* in the OC (Miwa et al., 2008; Müller et al., 2006). The absence of WUS activity leads to SAM arrest early in development (Laux et al., 1996), whereas the loss of the *CLV* genes results in a vast enlargement of the SAM (Clark et al., 1993; Clark et al., 1995; Kayes and Clark, 1998) through a phenomenon called fasciation. Several instances of fasciated tissues have been observed in the wild, as well as in crop plants, including beef tomatoes and the maize *fasciated ear2* mutant (Taguchi-Shiobara, 2001). It is thought that fasciation is caused by stem cell overproliferation in the CZ (Brand et al., 2000; Busch et al., 2010; Dao et al., 2022; Kwon et al., 2005; Lenhard and Laux, 2003; Müller et al., 2008; Nimchuk et al., 2011; Whitewoods et al., 2018; Wu et al., 2005), or conceivably, by lower rates of cell transit from the CZ to the PZ (Laufs et al., 1998).

Over the years, *CLV3* has been the only known genetic marker to study stem cells at the shoot apex (Müller et al., 2006; Reddy and Meyerowitz, 2005), including for analysing *clv* mutants. While microarray analyses have uncovered several other genes that are enriched in the CZ (Aggarwal et al., 2010; Busch et al., 2010; Yadav et al., 2009), none have been extensively characterised. More recently, atomic force microscopy (AFM) experiments, which measure cellular mechanical properties, revealed that *CLV3*-expressing cells are stiffer than surrounding PZ cells, and that the onset of *CLV3* expression in flowers coincides with an increase in cell wall stiffness (Milani et al., 2014). Thus, although stem cell identity is both spatially and temporally associated with increased cell stiffness (Milani et al., 2014), it is unknown how these mechanical patterns are altered when stem cell regulation is perturbed.

The core stem cell regulatory network is also influenced by hormone signalling, including cytokinin and auxin (Gordon et al., 2009; Busch et al., 2010). Exogenous cytokinin treatment phenocopies the *clv* mutant by increasing *WUS* expression and decreasing *CLV1* expression (Lindsay et al., 2006). While exogenous auxin treatment induces organogenesis markers in the PZ, the CZ itself remains unaffected, displaying reduced responsivity to auxin (de Reuille et al., 2006). Whereas in the PZ, the auxin response factor MONOPTEROS (MP) activates the expression of lateral organ identity genes in an auxin-dependent manner (Berleth and Jürgens, 1993; Bhatia et al., 2016; Hardtke and Berleth, 1998; Yamaguchi et al., 2013), it also functions to repress CZ genes that are themselves activators of *CLV3* expression (Luo et al., 2018; Zhao et al., 2010). Despite these advances, it remains unclear how auxin signalling is perturbed in *clv* mutants, and furthermore, whether this plays a direct role in generating the fasciated *clv* phenotype.

In this study, we use quantitative approaches to study the effects of the loss of proper stem cell regulation that reveal hitherto unobserved cellular and tissular phenotypes, such as cell size and local surface curvature in *clv* mutant SAM. Analytical mechanical models predict that heterogeneities in stiffness and/or growth regimes are sufficient to account for the changes in surface curvature observed in the mutant. We provide experimental support for these predictions by showing that both growth and mechanical properties display regional differences, which are likely caused by the chimeric identities of mutant meristematic cells. Finally, auxin-response quantifications in *clv* meristems reveal a behaviour similar to undifferentiated WT PZ cells, rather than to stem cells.

## Results

### Altered cellular and tissular properties characterise fasciated *clv* SAM

In order to better understand the defects associated with abnormal stem cell regulation, we first quantified the highly fasciated phenotype of the canonical *clv3-2* allele (Clark et al., 1995). WT shoot meristems displayed a stereotypical dome shape, whereas *clv3-2* meristems comprised a large centrally-located bulge and two or more arm-like outgrowths that elongate laterally (Fig. 1A, B, Fig. S1A-C). Further analysis revealed that the central areas were usually devoid of any discernible cellular organisation, whereas the lateral arms were often composed of cell files (Fig. S1D, E). These results were consistent across three *clv1* and four *clv3* alleles (Fig. S1A-C) in our growth conditions, although individual plants showed varying degrees of fasciation.

**Figure 1.**
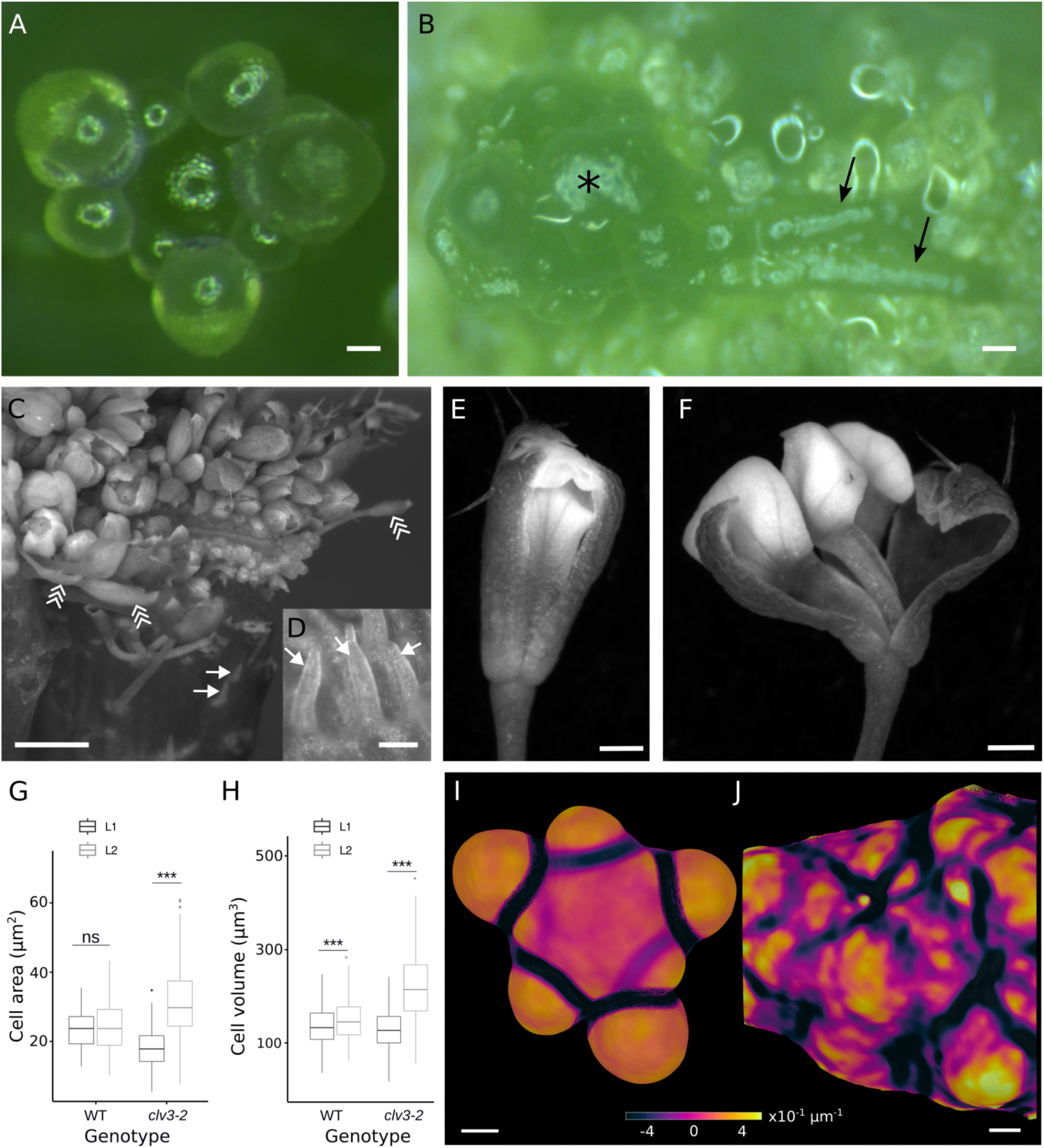
Altered cell and tissue properties characterise fasciated SAM. **(A, B)** Bright field images showing dissected WT L*er* (A) and *clv3-2* (B) SAM. The asterisk in (B) shows the central outgrowth and arrows point out two linear elongations. **(C-F)** Bright field images showing a dissected *clv3-21* SAM with filaments (arrows) on the side of the stem and abnormally shaped flowers (arrowheads) (C), a close-up of the filaments (D), and *clv3-21* abnormally-shaped flowers (E, F). **(G, H)** Boxplots of cell areas (G) or cell volumes (H) in the L1 (black) and L2 (grey) cell layers of WT and *clv3-2* SAM. Mean ± s. d. areas (G) are 23.9 ± 5.8 µm^2^ (WT L1) and 24.5 ± 7.2 µm^2^ (WT L2); and 18.4 ± 5.3 µm^2^ (*clv3* L1) and 31.2 ± 9.8 µm^2^ (*clv3* L2). *P*-values (Welch’s t-test) = 0.545 (WT) and 4.65e-54 (*clv3-2*). n = 103 (L1) and 115 (L2) cells from 3 WT SAM; n = 400 (L1) and 244 (L2) cells from 3 *clv3-2* SAM. Mean ± s. d. volumes (H) are 136.5 ± 40.5 µm^3^ (WT L1) and 148.7 ± 44.5 µm^3^ (WT L2); and 129.5 ± 40.9 µm^3^ (*clv3* L1) and 249.1 ± 130.7 µm^3^ (*clv3* L2). *P*-values (Welch’s t-test) = 2.32e-9 (WT) and 1.53e-204 (c*lv3-2*). n = 1468 (L1) and 529 (L2) cells from 4 WT SAM; 6426 (L1) and 3428 (L2) cells from 3 *clv3-2* SAM. **(I, J)** Colour maps quantifying minimal curvature in representative WT Col-0 (I) and in *clv3-2* (J) SAM. The *clv3-2* SAM (J) is only a small region of the entire meristem. Colourmap: warm helix (I, J). Scale bars: 30 μm (A, B, I, J), 1 mm (C), 200 μm (D), 100 μm (E, F).

We also detected novel floral phenotypes. First, we frequently observed radially symmetrical, filament-like organs either growing on the flanks of the meristem or within the inflorescence itself (Fig. 1C, D). Such filaments were recently noted as aborted flowers in double mutants of *clv2* or *clv3* and with auxin biosynthesis mutants (John et al., 2023; Jones et al., 2020). Secondly, while the majority (>90%) of flowers showed previously described phenotypes of supernumerary organs (Clark et al., 1993; Clark et al., 1995), the rest displayed two types of opposing phenotypes. One group of flowers bore two or three sepals and petals, a single stamen and no carpels (Fig. 1C, E, F), reminiscent of the inner-whorl phenotypes of *wus* mutants (Laux et al., 1996), whereas a second group displayed a tubular phenotype with fused sepals (Fig. 1C). These flowers were distinctly smaller than those with supernumerary organs. These novel floral phenotypes were observed in all *clv1* and *clv3* alleles.

Because it had been suggested that cell size in *clv* mutants differs from the WT (Laufs et al., 1998), we wondered whether we could use such changes in cellular morphologies to further characterise the mutant phenotypes described above. We quantified cell size in two ways: firstly, we calculated cell areas by acquiring images of meristems from WT and multiple *clv1* and *clv3* SAM, and extracting slices near the midpoints of cells in the L1 and L2 layers, which we then segmented using the MorphoGraphX software (Barbier de Reuille et al., 2015). In WT SAM, the average area was similar in both cell layers, with L1 cells at 23.9 ± 5.8 µm^2^ and L2 cells at 24.5 ± 7.2 µm^2^ (Fig. 1G). In contrast, *clv3-2* mutants exhibited significant differences in the L1 and L2, with L2 cells on average 69% larger than L1 cells – 31.2 ± 9.8 µm^2^ in the L2 versus 18.4 ± 5.3 µm^2^ in the L1 (Fig. 1G). Secondly, we measured cell volumes in WT and *clv* SAM, by generating 3D reconstructions using the MARS pipeline (Fernandez et al., 2010), which involves imaging samples from multiple angles and fusing those images to generate high-resolution images that are then segmented in 3D (Fig. S1F). Consistent with our findings for cell areas, we found that while L1 and L2 cells had comparable volumes in the WT (136.5 ± 40.5 µm^3^ and 148.7 ± 44.6 µm^3^, respectively), L2 cells in *clv3-2* SAM were almost twice as large as L1 cells (249.1 ± 130.7 µm^3^ vs 129.5 ± 40.9 µm^3^) (Fig. 1H). Similar trends were observed for cell size in other *clv1* and *clv3* alleles, suggesting that altered cell morphology is a general feature of these mutants (Fig. S1G).

We had noticed that meristems in *clv3* mutants are less smooth than WT meristems. We quantified this by measuring curvature at the sample surface, using a method that detects the outer surface in MARS-reconstructed 3D meristems (Kiss et al., 2017). We used MorphoGraphX to calculate curvature within a 10-µm radius of every pixel on the surface. In WT plants, the meristem dome and young flowers were regions of uniformly positive curvature, separated by clearly defined boundary areas with negative curvature (Fig. 1I). However, the imaged areas of *clv1* and *clv3* SAM, which are distant from the regions where flowers form, displayed a much more variable surface curvature, with crests and troughs of varying shapes and depths distributed throughout (Fig. 1J). Together, our data reveal that *clv* mutants exhibit abnormal cellular traits and buckling defects that contrast with the stereotypical characteristics of WT meristems.

### Morphoelastic models reveal how growth and stiffness contribute to tissue buckling

To elucidate a theoretical basis for the buckling defects of *clv* mutants, we next generated analytical models of SAM development using a growth and remodelling framework (Goriely, 2017). Specifically, we adapted a model developed to explore the biomechanical basis of morphogenesis in unrelated systems that display similar buckling behaviour to that observed in *clv* mutants (Almet et al., 2019; Moulton et al., 2013). We modelled the SAM as an axially growing planar elastic rod (the outer L1 layer), attached to an elastic foundation (the inner layers), and clamped at the two ends (Fig. S2). We used the theoretical model to explore how growth stretch, a proxy for tissue growth, and/or mechanical properties, a proxy for cellular stiffness, might contribute to the altered morphology of *clv* mutants (see Supplementary Materials and Methods).

First, we used linear stability analysis to plot the output shapes when either the growth stretch (γ) alone or the system stiffness (*k̂*, the ratio of foundation and rod stiffnesses) was varied. These analyses showed that when the foundation and rod had similar mechanical properties (*k̂* = 1) and when growth stretch was at a low, subcritical value (*γ*\*_inf_= 5.82), the output shape resembled the smooth dome of a WT meristem (Fig. 2A). In contrast, when the growth stretch was raised to a near-critical point (*γ*\*_inf_= 8.99), the size of the dome increased, while remaining smooth. A further increase in growth stretch generated various modes of buckling (Fig. 2A) that broadly capture the curvature defects of *clv* mutants. Next, we investigated the role of system stiffness (*k̂*) on SAM morphology, by varying stiffness within a background of constant growth stretch (*γ*\* = 12.23) (Fig. 2B). Whereas no buckling was visible for higher values of stiffness (*k̂* ≥ 1), lowering it (*k̂* = 0.5) led to the appearance of buckling in the rod (Fig. 2B), which was similar to those obtained using growth values above the critical point (Fig. 2A). Taken together, these analytical results indicate that in a biological context, cellular growth and stiffness characteristics could influence tissue architecture of the SAM.

**Figure 2.**
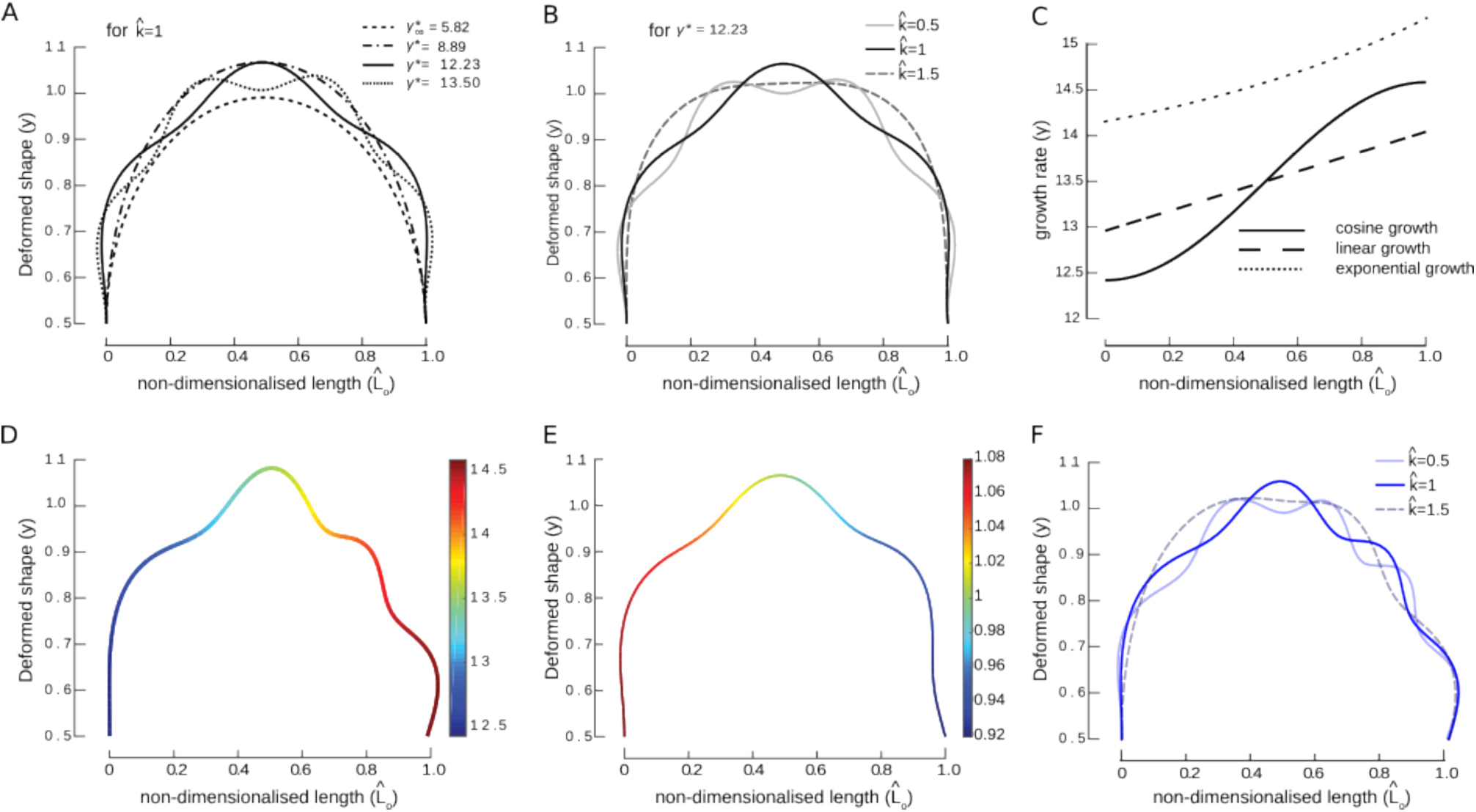
Growth and stiffness influence SAM shapes via growth-induced buckling. **(A)** Deformed shapes due to growth-induced buckling obtained for different values of critical growth rates (γ*) causing various buckling modes, with the stiffness of the underlying foundation fixed (*k̂* = 1). **(B)** Deformed shapes due to stiffness-induced buckling obtained for different values of foundation stiffness (*k̂* = 0.5, *k̂* = 1, *k̂* = 1.5), with the growth rate fixed (γ* = 12.23). **(C)** The choice of growth law affects the deformed shapes. We select three different forms of growth laws for the same value of growth rate *γ*_0_: cosine, linear and exponential. **(D)** Deformed shape obtained as a result of growth-induced buckling in the case of cosine growth distribution. **(E)** Deformed shape obtained as a result of stiffness-induced buckling in the case of cosine stiffness distribution. **(F)** Effect of foundation stiffness on the asymmetric rod shapes obtained using growth heterogeneity.

In contrast to the symmetrical shapes obtained above, *clv* mutant SAM in fact display highly variable buckling, such that the positions, numbers and amplitudes of folds differ within, and between, individual mutant meristems. We reasoned that this variability might result from local differences in growth, stiffness or both. To investigate the effects of such heterogeneities on buckling, we replaced constant growth stretch or system stiffness with one of three distinct functions to generate differential distributions across the rod (Fig. 2C). Indeed, all three functions generate variable buckling over the length of the rod when either growth stretch (Fig. 2D, Fig. S3B-B’) or system stiffness (Fig. 2E) is spatially distributed, with the resulting shapes often closely resembling the asymmetry and variable buckling amplitudes visible in sections through various *clv* SAM in the literature. These results suggest that local differences in growth or stiffness within the SAM could explain the phenotypes of *clv* mutants.

Finally, we asked whether variations in system stiffness and growth stretch could act together to generate even greater differences in buckling characteristics. To address this question, we used a cosine spatial growth stretch distribution and examined how buckling was affected when system stiffness (*k̂*) was varied between 0.5 and 1.5. Indeed, for identical distributions of growth, rod buckling varied locally as a function of stiffness (Fig. 2F); these analyses suggest that small changes at the cellular level could alter shape locally.

Taken together, our results indicate that growth and mechanical traits are sufficient to describe the differences in meristem shape between WT and *clv* SAM. Furthermore, local variations of these two cellular parameters could lead to the local curvature defects observed in *clv* meristems.

### Heterogeneous growth correlates with buckling in *clv* mutants

We tested predictions from the analytical model by first examining whether *clv* mutants display local heterogeneities in growth and, more specifically, whether these differences correlate with areas of differential curvature. We did this by dissecting WT and *clv3-7* meristems, placing them on culture medium and imaging them once a day over 72 hours by confocal microscopy. We then segmented the image stacks, tracked cell lineages, and analysed cell size, growth and curvature over the tissue (Fig. 3A-H, Fig. S4A-C). In the WT, cells had a mean size of 22.8 ± 5.8 µm^2^ at the start of the experiment and over 72 hours their daughters had grown 5.5 ± 1.3 fold (Fig. 3A-C). Of these, lineages that remained entirely within the meristem dome grew 4.9 ± 1.1 fold. In contrast, cells at the very top of the *clv3* meristem had an initial size of 17 ± 6.2 µm^2^ and had grown 1.5 ± 0.3 fold over 72 hours (Fig. 3E-G).

**Figure 3.**
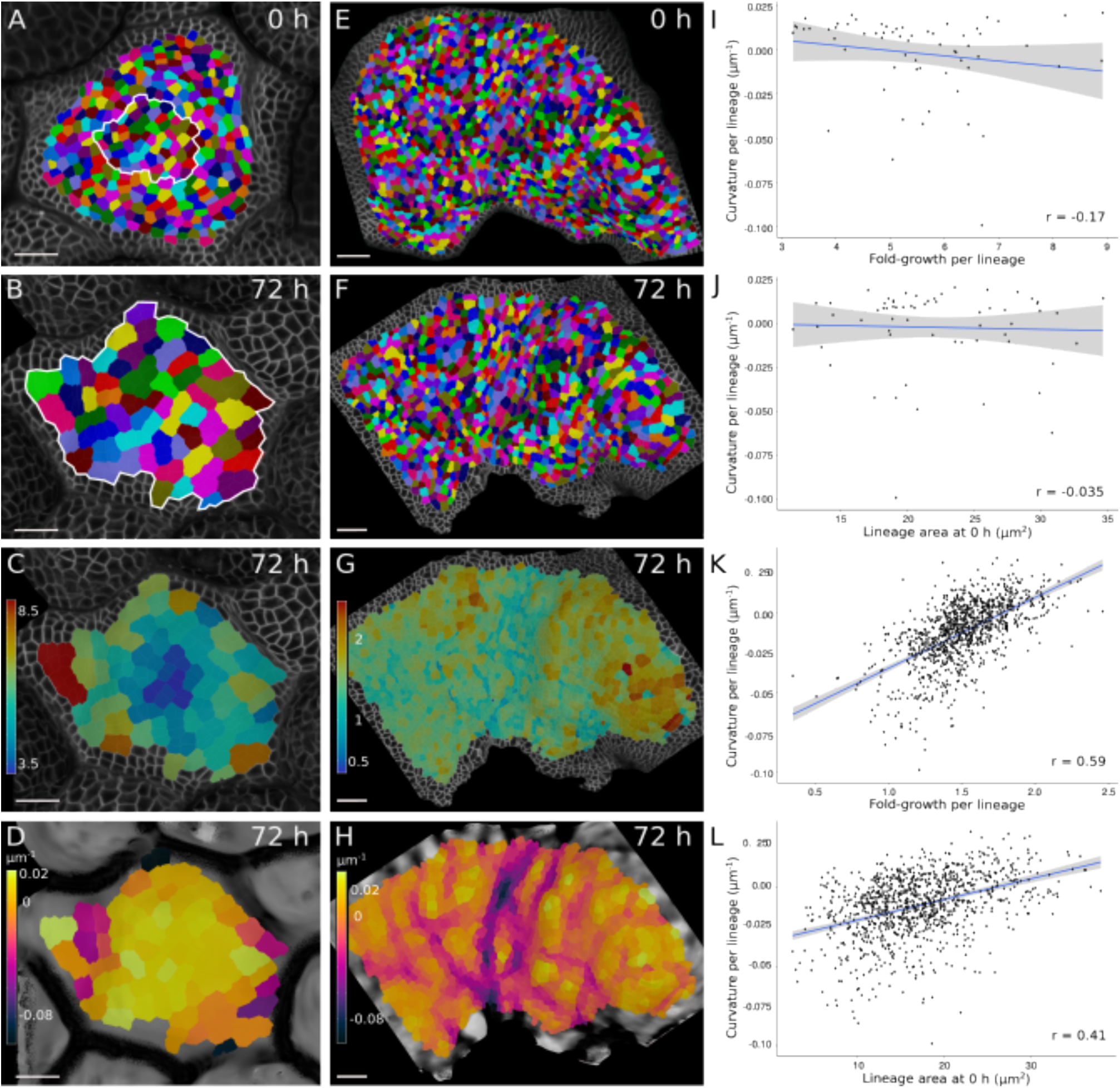
Regional growth heterogeneities correlate with surface buckling in *clv* mutants. **(A-H)** 3D surface reconstruction of a WT Col-0 (A-D) and a *clv3-7* (E-H) SAM, expressing the *pUBQ10::Lti6b-tdTomato* membrane marker, at the start of the experiment (A, E) and 72 h later (B, F). Solid white line in (A) and (B) delineates the same group of cells. Segmented cells are coloured by lineage and outlines of unsegmented cells are shown. For each lineage, heatmaps for either growth fold-change over 72 h (C, G) or minimal curvature (D, H) are represented on the second time point. **(I-L)** Scatter plots of minimal curvature per lineage as a function of fold-growth per lineage (I, K), or of minimal curvature per lineage as a function of initial lineage surface area (J, L) in WT (I, J) and *clv3-7* (K, L) SAM. Regression lines representing the best fit are shown in blue, and confidence intervals as shaded regions around them. n = 64 WT lineages (I, J); n = 1064 *clv3* lineages (K, L). Pearson’s correlation coefficients (r) are shown on the plots. *P*-values: *p* = 0.18 (I); *p* = 0.79 (J); *p* < 0.001 (K); *p* < 0.001 (L). Colourmap: Turbo (C, G); warm helix (D, H). Scale bars: 20 µm (A-H).

In the WT, a gradient of differential growth is observed from the centre to the periphery of the meristem, whereas in the mutant, groups of cells with varying growth rates are seemingly scattered across the enlarged meristem (Fig. 3C, G). To determine whether this observation is relevant in the context of buckling in *clv* meristems, we then measured curvature at the surface of WT and mutant samples, determined the mean value per cell (Fig. 3D, H, Fig. S4C), and tested correlations between cell size, growth rate and local curvature (Fig. 3I-L, Fig. S4D-E). WT meristems showed no significant correlations between any parameters (Pearson’s correlation coefficients (r) between: growth and curvature, *r* = -0.17, *p* = 0.18; size and curvature, *r* = -0.035, *p* = 0.79; growth and size, *r* = -0.23, *p* = 0.07) (Fig. 3I-J, Fig. S4D). *clv3* mutants, however, displayed significant moderate or strong positive correlations between all parameters (Pearson’s correlation coefficients between: growth and curvature, *r* = 0.59, *p* < 0.001; size and curvature, *r* = 0.41, *p* < 0.001; growth and size, *r* = 0.36, *p* < 0.001) (Fig. 3K-L, Fig. S4E). These analyses show that in *clv3* mutants, both cell size and growth rate are variable across the tissue and are linked to local buckling characteristics.

### Loss of *CLV* activity is associated with reduced and variable epidermal cell stiffness

In addition to growth heterogeneity, our analytical models predicted that buckling is also dependent on stiffness changes in the SAM. We had previously shown that *CLV3*-expressing CZ cells are stiffer than PZ cells (Milani et al., 2014). Because the *clv* phenotype is thought to be caused by stem cell overproliferation, we predicted that cells in *clv* meristems might bear higher values of stiffness. To test this, we probed epidermal cells in WT, *clv1* and *clv3* SAM using atomic force microscopy (AFM), as published (Milani et al., 2014; Beauzamy et al., 2015) (Fig. S5A-D). The resultant force-displacement curves at each measured point from individual AFM scans were analysed to determine the apparent Young’s modulus up to a depth of 100 nm, and these values were subsequently corrected to remove artefacts caused by local slope (Fig. 4A-D). The data were either pooled and plotted as density curves (Fig. 4E, F) or analysed by individual scans as box plots (Fig. 4G, H). When compared to the appropriate WT accession, we found that both *clv3-2* (Fig. 4E, G) and *clv1-8* cells (Fig. 4F, G) were significantly softer, with apparent Young’s moduli that were 15-20 % lower than WT cells (Fig. 4G, Fig. S5E, Table S1, Table S2).

**Figure 4.**
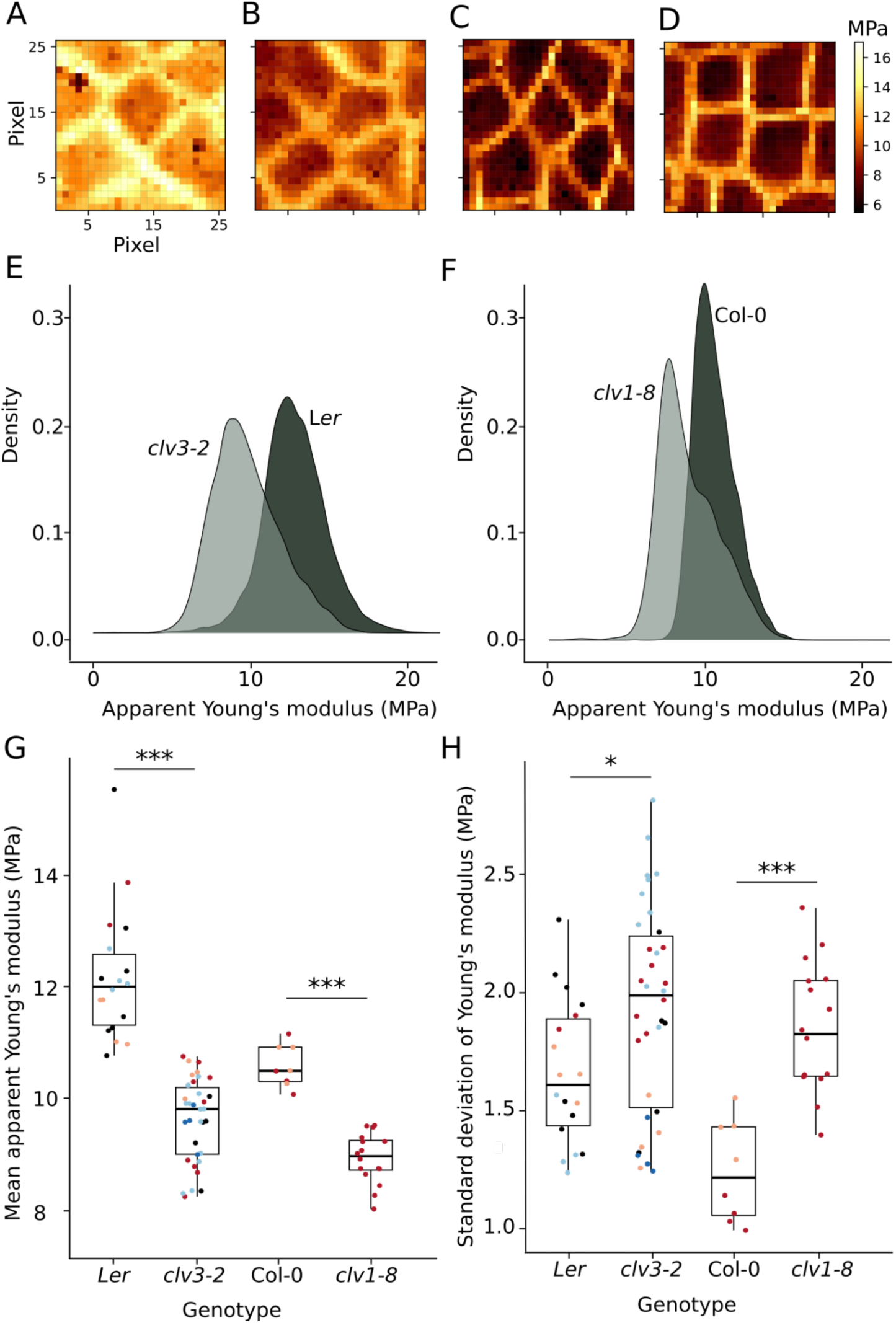
Loss of *clv* is associated with reduced and heterogenous stiffness. **(A-D)** Maps showing apparent Young’s moduli derived from atomic force microscope scans of representative L*er* (A), Col-0 (B), *clv3-2* (C), and *clv1-8* (D) SAM. **(E, F)** Density plots of apparent Young’s moduli from all probed WT (L*er*) and *clv3-2* (E), and WT (Col-0) and *clv1-8* (F) samples. **(G-H)** Box plots of either mean values (G) or standard deviation values (H) of apparent Young’s moduli measurements of scans from all probed samples. Black dots show values of different samples probed by a single scan, whereas coloured dots represent values from samples bearing multiple non-overlapping scans, with each colour identifying an individual sample. Overall mean values derived from individual mean apparent Young’s moduli (G): L*er*, 12.2 ± 1.2 MPa; *clv3-2*, 9.7 ± 0.7; Col-0, 10.58 ± 0.4; and *clv1-8*, 8.94 ± 0.5. Overall mean values derived from individual standard deviations (H): L*er*, 1.7 ± 0.3 MPa; *clv3-2*, 1.9 ± 0.4; Col-0, 1.2 ± 0.2; *clv1-8*, 1.9 ± 0.3. *P*-values (Welch’s t-test): WT–*clv3*, 6.65e-8 and WT–*clv1*, 1.85e-7 (G); WT–*clv3*, 3.4e-2 and WT–*clv1*, 7.1e-5 (H). n (non-overlapping scans/independent meristems): L*er*, 18/11; Col-0, 8/2; *clv3-2*, 34/8; and *clv1-8*, 16/1.

While these results show that *clv* mutant SAM bear different physical characteristics to WT cells, they do not reveal whether they also display greater heterogeneity. To test this, we plotted standard deviation values for each scanned region by genotype (Fig. 4H) and found that *clv* mutant samples indeed display a significantly greater variability in values of apparent Young’s moduli when compared to the WT. This altered regime is also visible in the density plots (Fig. 4E, F), where mutant distributions are notably broader than in the WT. Furthermore, scatter plots of the dataset show that mean and standard deviation are not correlated (Fig. S5F-I). Consistent with our analytical models, these results suggest that altered local stiffness and increased heterogeneity could indeed contribute to tissue buckling in *clv* mutants.

### Genetic domain separation is lost in *clv* SAM

We reasoned that the local heterogeneities in growth and stiffness described above might be caused by underlying variations in genetic identity. To examine this, we first retested the expression patterns of OC and CZ markers using mRNA *in situ* hybridisation assays in WT and different *clv3* alleles. We used *WUS* to examine OC identity (Laux et al., 1996; Mayer et al., 1998), whereas for the CZ, we tested several genes previously identified as enriched in *CLV3*-expressing cells (Yadav et al., 2009) and retained the *APUM10* gene, which encodes a member of the Puf family, with a conserved Pumillio homology domain (Tam et al., 2010). *APUM10* was observed in a few centrally-located cells of the L1 and L2 in WT SAM (Fig. 5A), and *WUS* in an underlying group of cells (Fig. 5B). In *clv3* mutants, we detected both *APUM10* and *WUS* in broad domains occupying almost the entire meristem, with *APUM10* expressed principally in the L2 layer (Fig. 5C, C’) and *WUS* in deeper layers (Fig. 5D-D’). However, both genes were expressed in a discontinuous manner, with no detectable expression in scattered groups of cells. An inspection of published expression data in *clv* mutants revealed that such patchy expression was frequently visible, but had gone unremarked (Brand et al., 2000; Kinoshita et al., 2015; Schuster et al., 2014).

**Figure 5.**
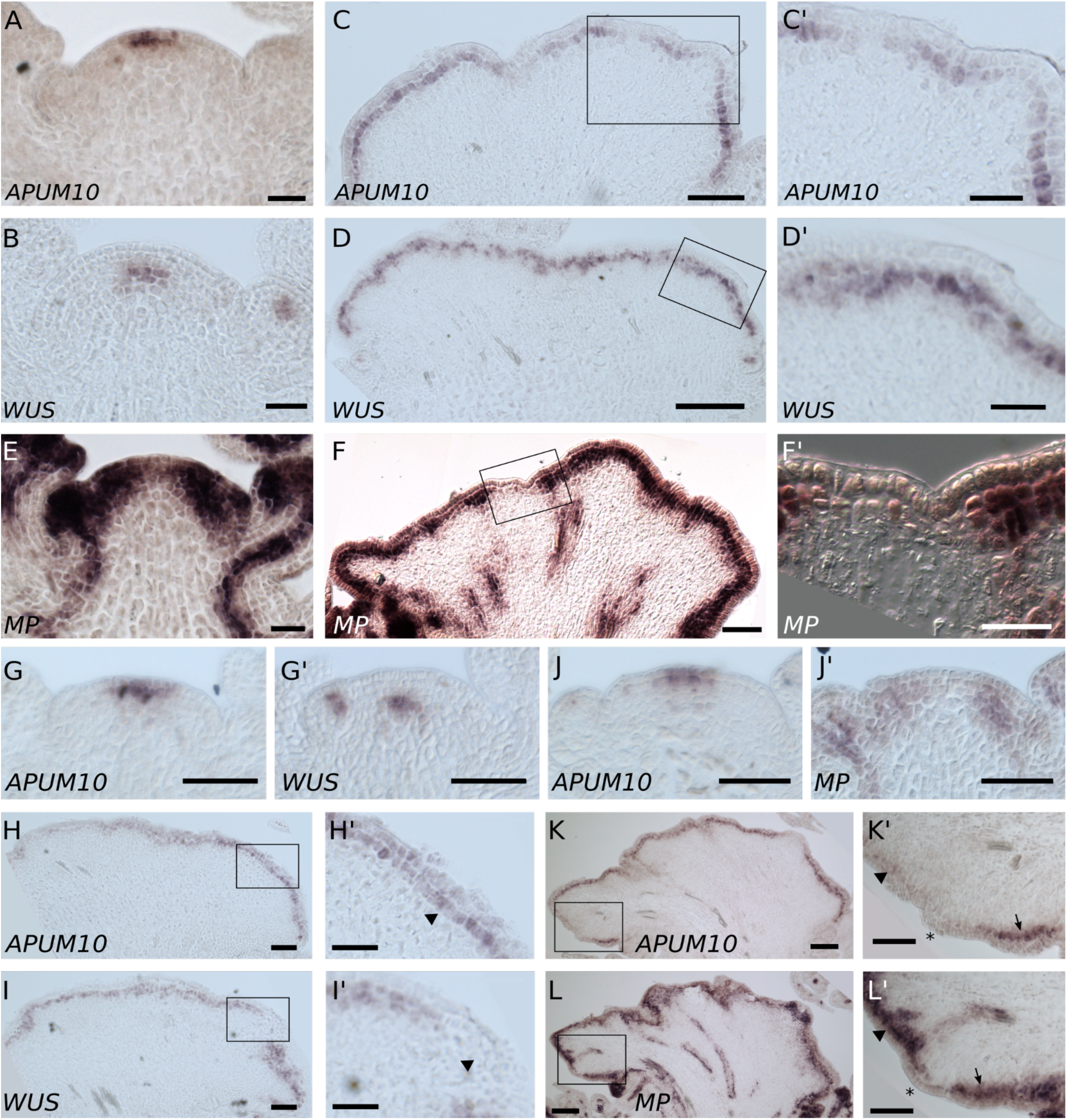
Gene expression is heterogenous in *clv3* SAM. **(A-F’)** Pattern of mRNA localisation for *APUM10* (A, C, C’), *WUS* (B, D, D’), and *MP* (E-F’) in WT L*er* (A, B, E) or *clv3-2* (C, C’, D, D’, F, F’) SAM. (C’, D’, and F’) are magnified views of the boxed regions in (C, D, and F), respectively. **(G-L’)** Pairwise localisation of *APUM10* (G, J, H, H’, K, K’), *WUS* (G’, I, I’), and *MP* (J’, L, L’) transcripts on alternating sections of meristems from individual WT L*er* (G, G’, J, J’) or *clv3-2* (H-I’, K-L’) plants. Arrowheads (H’,I’, K’, L’) indicate zones where only one of the two transcripts is detected, asterisks (K’, L’) indicate regions where neither is detected, and arrows indicate areas where both are detected. (H’, I’, K’, and L’) are magnified views of the boxed regions shown in (H, I, K, and L), respectively. Section thickness: 10 μm. Scale bars: 50 µm (A-C, E, F, G, G’, H’, I’, J, J’, K’, L’); 100 µm (D, H, I, K, L); 25 µm (C’, D’, F’).

One explanation for why some patches of cells in *clv3* mutants expressed neither CZ nor OC identity was that they might instead bear other identities. To test this, we localised mRNA for the *MP* gene (Hardtke and Berleth, 1998), which is expressed mainly in the WT PZ (Zhao et al., 2010) (Fig. 5E). In *clv* SAM, the PZ is thought to be restricted to areas at the base of the enlarged dome, in the region where flowers form. However, we found that in *clv3* mutants, *MP* is expressed in a broad and patchy manner (Fig. 5F-F’), similarly to the *APUM10* and *WUS* patterns. Taken together, our results suggest that identities of the three meristematic zones are present in overlapping groups of cells in *clv* apices.

We thought it possible that normal domain organisation might exist at a more local level. In order to investigate the positions of the OC and PZ markers relative to the CZ marker within *clv* SAM, we performed pairwise *in situ* hybridisations using alternating sections from individual fixed meristems, with one probe used on even-numbered sections, and a second on odd-numbered sections. In WT meristems, the *APUM10*, *WUS* and *MP* expression domains were never detected in similar regions within adjacent sections (Fig. 5G, G’, J, J’). However in *clv3* mutants, *APUM10* and *WUS* expression overlapped across the entire SAM (Fig. 5H-I’, Fig. S6), except for a few areas where *APUM10* was observed but *WUS* was not (Fig. 5H’, I’) and rare areas where *WUS* was expressed without *APUM10*. Similarly in *clv3* mutants, *APUM10* and *MP* were expressed in overlapping patterns, except for a few areas where *MP* was expressed without *APUM10* (Fig. 5K-L’, Fig. S7) and no instances where *APUM10* was expressed without *MP*. We also observed a few cell patches devoid of any signal.

Together, these results show that *clv3* meristems comprise a heterogeneous group of cells with mixed genetic identities, rather than a homogeneous population of stem cells. Furthermore, we observed similar expression patterns in SAM from other alleles of *clv1* and *clv3* (Fig. S8), suggesting that chimeric cell identities are a general feature of fasciated *clv* meristems.

### Exogenous auxin elicits a strong response in *clv3* meristems

Because cells in fasciated *clv3* SAM displayed traits not usually associated with stem cells, such as chimeric cell identities and reduced stiffness, we wished to better understand if they were also functionally different from stem cells. We chose auxin response as a functional readout for the following reasons: firstly, auxin-associated genes are differentially expressed in WT and *clv3* mutants (John et al., 2023). Secondly, a key aspect of organogenesis at the shoot apex is the presence of a CZ that is refractive to auxin signalling (de Reuille et al., 2006; Douady and Couder, 1996; Galvan-Ampudia et al., 2020; Vernoux et al., 2011), suggesting that auxin insensitivity is an indicator of stem cell identity. Thirdly, MP, a polarity regulator of the auxin efflux transporter PIN1, which itself promotes auxin accumulation and drives organogenesis (Bhatia et al., 2016), is misregulated in *clv* mutants (Fig. 5E, F).

In order to test how *clv* mutants respond to auxin, we dissected shoot apices from WT, *clv1* and *clv3* plants carrying the auxin signalling reporter, *pDR5rev::GFPer*, and treated them with exogenous auxin as described (Galvan-Ampudia et al., 2020), while monitoring expression every 24 hours. As is well understood, in 100 % of WT samples prior to treatment, GFP was localised in small groups of cells corresponding to the positions of floral anlagen, accompanied by low expression in the geometric centre of the SAM (Fig. 6A). 24 hours later, we observed high levels of GFP throughout the periphery of the SAM and in internal layers, whereas expression levels at the centre remained low (Fig. 6B). *clv* meristems differed from the WT in that prior to auxin treatment, we detected only very basal levels – several thousand-fold lower than in the WT – of GFP (Fig. 6C), whereas 24 hours after treatment, a clear response was visible in all samples (Fig. 6D). More specifically, a minority of meristems (1 out of 11 *clv3* SAM) showed high levels of GFP in both outer and inner layers (Fig. S9A, B), whereas in all other samples (10 out of 11 *clv3*, and 5 out of 5 *clv1*) expression levels were variable and patchy in outer layers, and high in inner layers (Fig. 6D).

**Figure 6.**
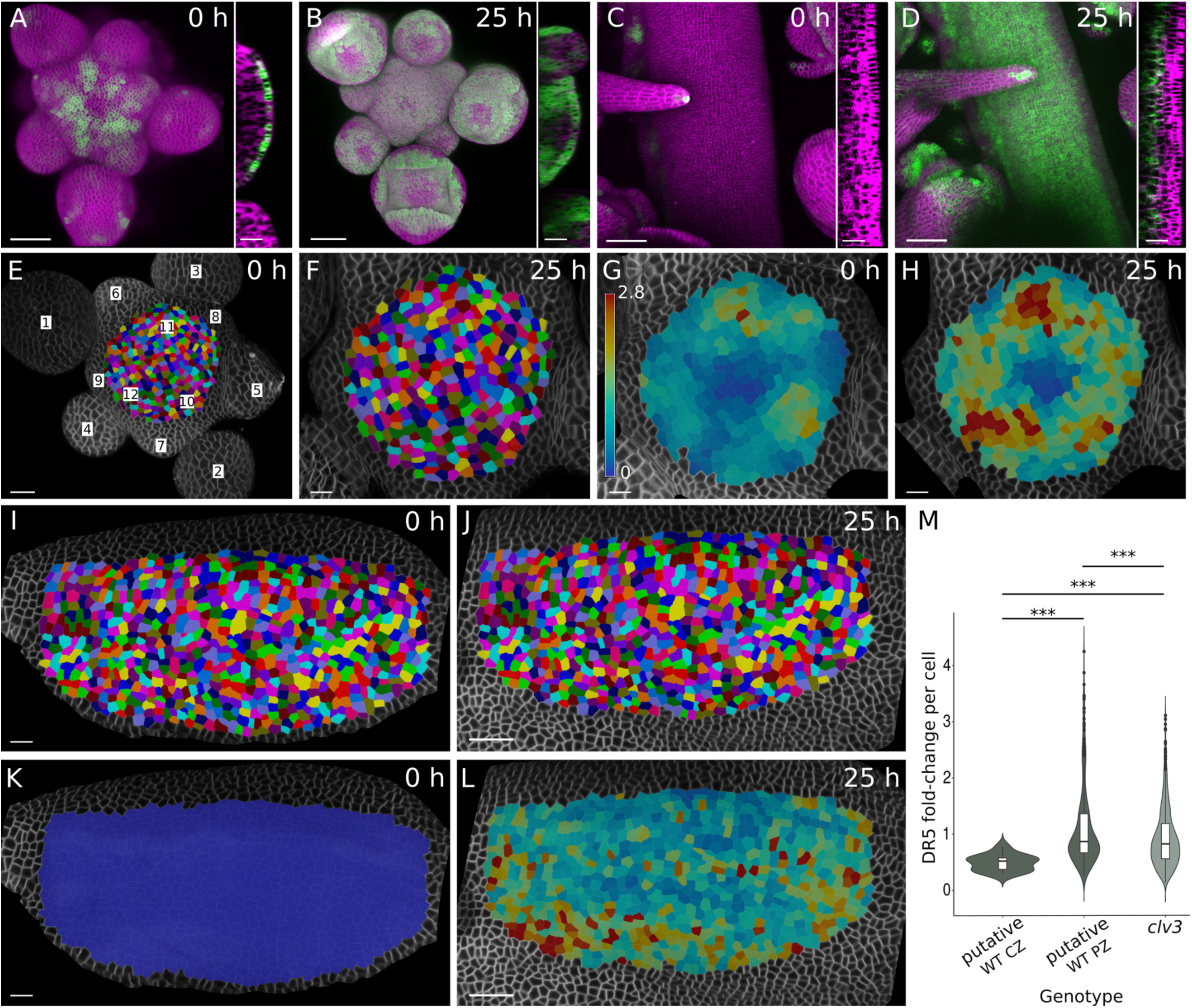
*clv3* meristematic cells respond strongly to exogenous auxin treatment. **(A-D)** Projections of confocal stacks showing the effect of IAA on WT Col-0 (A, B) and *clv3-2/clv3-21* (C, D) SAM expressing the *pUBQ10::Lti6b-tdTomato* and *pDR5::GFPer* reporters. Samples are shown either before (A, C) or 25 h after (B, D) a 5-hour IAA treatment. Orthogonal views of the SAM are shown alongside each panel. n = 12 WT SAM (5 independent experiments), n = 11 *clv3* SAM (4 independent experiments). **(E-L)** Quantification of response to IAA treatment observed in representative SAM from WT Col-0 (E-H) or *clv3-2/clv3-21* heterozygotes (I-L) carrying the *pUBQ10::Lti6b-tdTomato* and *pDR5::GFPer* reporters. Views of the SAM surface before (E, I) and after (F, J) treatment showing segmented cells coloured by lineage and unsegmented cells as grey outlines. Numbers in (E) represent the phyllotactic order of flowers from oldest to youngest. Heatmaps show DR5 signal per cell presented as a concentration (G, H, K, L) with a common colour scale (G). **(M)** Violin plot reflecting distributions of DR5 fold-change per cell lineage between *clv3* SAM and putative WT CZ and PZ (shown in E-L). *P*-values < 0.001 (Welch’s t-test). n = 12 cells for WT CZ; n = 300 cells for WT PZ; n = 713 cells for *clv3*. Colourmap: Turbo (G, H, K, L). Scale bars: 50 µm (top views in A-D); 20 µm (orthogonal sections in A-D, F-H, J, L); 10 μm (E, I, K).

We next quantified this response by segmenting image stacks of WT (Fig. 6E, F) or mutant (Fig. 6I, J) meristems before and after auxin treatment, and extracted average fluorescence values per cell (Fig. 6G, H, K, L). For each cell, we then calculated the ratio of expression between the two time points, normalised this across the sample, and compared values in the mutant to values for cells in either the geometric centre or the periphery of WT meristems. A histogram of cell signal intensities shows that before auxin treatment, 22 % of WT cells and 100 % of mutant cells were in a low-expressing category of signal intensities under an arbitrary threshold set to 1/10th of the highest value (Fig. S9C). After treatment with auxin, only 6.4 % of WT and 5.9 % of mutant cells remained in this category, suggesting that the *clv* meristem responded to exogenous auxin in a similar manner to the WT. Furthermore, plots of normalised intensity change per lineage show that cells from the putative WT CZ and WT PZ, as well as from the *clv3* mutant had significantly different distributions from each other, although qualitatively the mutant resembled the PZ more than the CZ (Fig. 6M). Similar results are obtained for *clv1* mutants carrying the *DR5* reporter (Fig. S9D, E). Taken together, these results show that cells in *clv* mutant SAM are capable of responding to auxin signalling, an output more associated with the WT PZ, where organ specification occurs.

## Discussion

The development of fasciated meristems due to the loss of *CLV* activity in *Arabidopsis* shoot apical stem cells has generally been interpreted as being caused by a massive increase in stem cell numbers (Brand et al., 2002; Busch et al., 2010; Dao et al., 2022; Kwon et al., 2005; Lenhard and Laux, 2003; Müller et al., 2006; Nimchuk et al., 2011; Whitewoods et al., 2018; Wu et al., 2005). Direct experimental evidence for this comes principally from a *CLV3* transcriptional reporter, which is expressed throughout the enlarged meristem of *clv* mutants (Brand et al., 2002; Reddy and Meyerowitz, 2005). However, because WUS protein is known to directly activate *CLV3* transcription (Yadav et al., 2011), and because *WUS* is expressed throughout *clv* meristems (Schoof et al., 2000), it is not surprising that *CLV3* reporter expression is broad in *clv* mutant. We suggest that the *CLV3* reporter might thus not be an appropriate marker to study cell identity in *clv* meristems. Furthermore, the use of the *CLV3* promoter for various misexpression studies of the role of stem cells in meristem maintenance might be inappropriate.

An alternate way of deciphering regional identities is by characterising cellular properties, which we have done using recently-developed approaches to precisely quantify cell volumes, areas and surface curvatures. Although we show that epidermal and subepidermal cell sizes in *clv* meristems are different from the WT, the effect of such changes on meristem structure is difficult to estimate. One possibility is that they generate a higher frequency of cell boundary overlaps between the L1 and L2 layers, which could generate a mechanically fragile structure, akin to a wall built with smaller cells in one layer and much larger bricks in the next.

Our analytical model attempts to explain meristem morphology through a mechanical framework, in contrast to various other modelling studies, which have shed light on different aspects of meristem shape and function, such as the role of auxin in phyllotactic patterning or homeostasis of the CLV-WUS feedback loop (Banwarth-Kuhn et al., 2022; Battogtokh and Tyson, 2022; Chickarmane et al., 2012; Fujita et al., 2011; Galvan-Ampudia et al., 2020; Gruel et al., 2016; Heisler et al., 2005; Jönsson et al., 2005; Jönsson et al., 2006; Klawe et al., 2020; Michael et al., 2023; Plong et al., 2021). Our results suggest that the specific buckling-like patterns displayed by *clv* mutants can be explained by a combination of local heterogeneities in growth rates and in mechanical properties. Indeed, we show that mutant tissues are 15-20% less stiff than the WT and display a wider range of stiffnesses. We had expected *clv* cells to be stiffer than WT cells, because *CLV3*-expressing WT CZ cells are stiffer than PZ cells (Milani et al., 2014), and because the prevailing view in the literature is that cells in *clv* SAM bear stem cell identity. Our findings indicate that this view might be incorrect.

A wavy surface can arise from buckling with the epidermis growing more than subepidermal layers, or from differences along the epidermis, with accelerated growth in bumps or growth constriction in the valleys. The results presented here cannot fully distinguish between these scenarios. However, a recent study revealed that cellular turgor pressure is anticorrelated to the size and number of neighbours of a given cell, but is correlated to its growth rate, such that smaller cells with fewer neighbours display higher pressure and higher growth (Long et al., 2020). In this context, our results showing a large epidermal-to-subepidermal cell size differential suggest that subepidermal cells in *clv* mutants are less stiff and grow less than the significantly smaller epidermal cells, which is consistent with predictions from our analytical model, that buckling occurs when inner layers are less stiff than the L1. A further experimental validation of this would require techniques to measure stiffness or pressure in internal layers.

That tissue-level shape robustness emerges from cell growth and cell size variability averaged through space and time has been shown in multiple contexts (Hervieux et al., 2016; Hong et al., 2016; Kamimoto et al., 2016; Tsugawa et al., 2017). Interestingly we find that overall, *clv* mutants grow at a much lower rate than the WT, indicating that the enlarged meristem of mature plants is likely to be a result of the amplification of early changes through time, rather than due to increased proliferation later in development. A recent study has shown that impairing cellulose synthesis decreases both cell proliferation and cell stiffness, leading to smaller SAM, and that in the context of a *clv1* mutant, it results in a dramatic reduction of the fasciated phenotype (Sampathkumar et al., 2019). Since we now show that *clv* mutants themselves have slower growth and reduced stiffness, it is not evident how a further reduction of both parameters could result in an amelioration of the *clv* phenotype. A recent study shows that cell expansion in parts of the meristem can be accounted for by stress-dependent orientation of cell division planes (Banwarth-Kuhn et al., 2022). Thus the early effects of lowered cellulose synthase activity in reducing tensile stress may prevent the *clv* phenotype from developing in plants with impaired cellulose synthase activity. Furthermore, an important distinction is that our study focuses on local buckling, rather than on global fasciation.

Mechanical and growth heterogeneities in *clv* mutants may be caused by underlying perturbations in genetic patterns. Variable and patchy expression of CZ and OC markers are in fact visible in the literature (Brand et al., 2000; Reddy and Meyerowitz, 2005; Schlegel et al., 2021; Schoof et al., 2000), although few studies have remarked upon, or interpreted, these data. It is unclear precisely how this pattern arises in the mutant and how it behaves over time. Furthermore, chimeric identities might exist in regions of fate transit within WT meristems, for instance at the CZ-PZ or the PZ-flower boundaries. In this context, it would be informative to look at the expression of a related negative feedback loop that functions specifically in the PZ (Schlegel et al., 2021).

In the absence of external stimuli, cells in *clv* meristems clearly do not undergo differentiation, like WT CZ cells. However, we show that cells in *clv* meristems are more akin to WT PZ cells in their capacity to respond to auxin signalling, perhaps due, at least in part, to their heterogeneous identities. Bhatia et al. (Bhatia et al., 2016) show that for organogenesis to occur, spatial differences in MP are a necessary condition. This is indeed the case in *clv* mutants, where *MP* expression is both broad and patchy, suggesting that improper auxin distributions or signalling contribute to the *clv* phenotype. It is thus unclear why organogenesis is affected in *clv* plants. One possibility is that CLV signalling might regulate auxin activity directly, such as in the moss (Nemec-Venza et al., 2022). Alternatively, because auxin accumulation in space and time is crucial for organogenesis (Galvan-Ampudia et al., 2020; Reddy and Meyerowitz, 2005; Reinhardt et al., 2000; Vernoux et al., 2000), it is possible that the abnormal shapes of *clv* SAM do not allow proper auxin patterns to be generated over most of the meristem. In this model, an initial change in the geometry of *clv* SAM would lead to perturbations in auxin fluxes, thereby inhibiting the production of lateral organs, which are a major auxin source (Shi et al., 2018). The presence of fewer primordia would generate even lower auxin levels in the SAM and lead to an increase in size (Shi et al., 2018). In support of this, the novel floral phenotypes we reveal are reminiscent of certain auxin pathway components (Cheng et al., 2006). Furthermore, it was recently shown that *clv* SAM are sensitised to auxin levels (John et al., 2023). A detailed analysis of auxin fluxes and auxin signalling in *clv* meristems will be necessary to understand the role of auxin in fasciation.

Taken together, all examined parameters in this study show that cells in *clv* SAM are different from WT CZ cells, and that they do not fit the current idea of typical stem cells. We suggest that stem cell identity needs to account for genetic, mechanical and functional parameters. Reporters for diverse cellular properties, as well as detailed real-time analyses, will help elucidate how chimeric identities arise and what their effects are on plant architecture.

## Materials and methods

### Plant material and culture conditions

The following plant lines were used in this study: L*er* and Col-0 as wild types, *clv3-2* (Clark et al., 1995), *clv3-7* (Fletcher et al., 1999), *clv1-8* (Medford et al., 1992), *pCLV3::GFPer* (Reddy and Meyerowitz, 2005), *pDR5rev::GFPer* (Friml et al., 2003), *pDR5::3xVENUS-N7* (Vernoux et al., 2011), and *pUBQ10::LTI6b-TdTomato* (Shapiro et al., 2015).

Seeds were sown on soil, and placed in short-day conditions (8 hrs light, 20°C, 50-60% humidity and 16 hrs dark, 16°C, 50-60% humidity) for ten days. Seedlings were then transplanted into individual pots and placed back into the short-day conditions growth chamber. One month after transplantation, plants were transferred from short-day to long-day conditions (16 hrs light period, 20°C, 60% humidity and 8 hrs dark period, 19°C, 60% humidity) until flowering, which is when we used plants for further experiments (in situ hybridisations, confocal microscopy, AFM). The light sources were LED fixtures (Valoya, C75, spectrum NS12), with an intensity of 150 μmol.m^-2^.s^-1^.

### CRISPR mutagenesis of CLV3

Generation of *CLV* mutants was carried out using a multiplexing CRISPR system (Stuttmann et al., 2021) to create the *clv3-21* allele in Col-0 plants carrying a *pUBQ10::LTI6b-tdTomato* reporter construct. Two guide RNAs targeting *CLV3* (5’-gATCTCACTCAAGCTCATGC-3’ and 5’-TCAAGGACTTTCCAACCGCA-3’) were designed using the Benchling software (www.benchling.com) and placed under the control of the *U6* promoter in the pDGE333 and pDGE334 ‘level 0’ shuttle vectors as described (Stuttmann et al., 2021). These sgRNA constructs were subsequently assembled in the CAS9-carrying pDGE347 binary vector, along with the FAST-red marker (fluorescence-accumulating seed technology) (Shimada et al., 2010) and the *bialophos resistance* gene (BAR) that confers resistance to glufosinate ammonium (Basta) for positive selection of transformants. T1 generation seeds that were positive for FAST-red were germinated on plates containing Basta and target genes from putative transformants were sequenced by Sanger sequencing, screened for *clv*-like phenotypes in the T2 and T3 generations and resequenced to generate independent homozygous lines. The *clv3-21* allele carries an insertion of an A after the second base pair of exon 2, leading to a frameshift at position 24 of the CLV3 protein.

### Bright field photos

Bright field photographs of plants were taken with a binocular Nikon SMZ18 equipped with an objective Nikon SHR Plan Apo 1X WD:60, and an ORCA-Flash4.OLT camera.

### Plant dissection and preparation for confocal microscopy

SAM were grown on soil and dissected soon after bolting. An approximately 1 cm region at the apex of the stem was placed in apex culture medium (ACM: ½ MS medium, 1% [w/v] sucrose, pH adjusted to 5.8 with 1M KOH, 0.8% [w/v] agarose, supplemented with 1X vitamins (1000X stock: 5 g myo-inositol, 0.05 g nicotinic acid, 640.05 g pyridoxine hydrochloride (B6), 0.5 g thiamine hydrochloride (B1), 0.1 g glycine, H_2_O to 50 mL, filter before aliquoting) and cytokinins (BAP; 125-175 nM final)), and flower buds were removed until the shoot meristem was sufficiently exposed for proper imaging. Dissected samples were placed in a growth cabinet under long-day conditions until confocal observation.

### Confocal imaging

Imaging was carried out on a Leica TCS SP8 (DM6000 CS) upright confocal laser scanning microscope, equipped with a 25x water dipping lens (Leica HC FLUOTAR L 25x/0.95 W VISIR) and a Leica HyD hybrid detector. Either plants expressed the *pUBQ10::LTI6b-TdTomato* fluorescent plasma membrane marker, or FM4-64 (ThermoFisher Scientific, T13320) was used to mark the plasma membrane as previously described (Fernandez et al., 2010). FM4-64 was excited with a 488 nm laser diode and detected at 600-640 nm. GFP was excited at 488 nm and detected at 500-520 nm. VENUS was excited at 514 nm and detected at 520-535 nm. tdTomato was excited at either 514 or 552 nm and detected at 560-600 nm.

### Image analysis

Fiji (Schindelin et al., 2012) was used to generate 2D projections from 3D confocal stacks (3D viewer plugin) and to generate orthogonal slices.

The MultiAngle Reconstruction and Segmentation (MARS) pipeline (Fernandez et al., 2010) was used to generate 3D reconstructions of confocal stacks. For this, shoot meristems were imaged from multiple angles by means of a custom-made device to tilt and/or rotate the sample by 15-20 degrees between acquisitions. After fusion using MARS, the external contours of the sample were detected using the Level Set Method (Kiss et al., 2017) and this contour was then used to segment the 3D reconstructed sample using a watershed segmentation algorithm. Cells were then extracted from the segmented image for the L1 and L2 tissue layers and the volume of each cell was calculated.

MorphoGraphX (Barbier de Reuille et al., 2015; Kiss et al., 2017) was used to measure cell size, growth and curvature for individual image stacks. For cell size, we determined the image slice within the L1 and L2 layers that was orthogonal to the axis of acquisition, and which contained the midline of the group of cells near the centre of the image (in order to avoid cells that had been imaged from the side). We then segmented and extracted cell areas. For growth, we first identified the tissue surface, generated the mesh, projected signal from the first cell layer (usually 5.5-6 μm), watershed-segmented the mesh, and determined cell lineages over two or more time points. Minimal curvature was calculated within a 10-μm radius for each point in the image, and for segmented tissues, was then normalised to the size of each cell. R was used to plot the data and run statistical tests.

### Statistical analysis

Plots and statistical analyses were done with R studio version 2023.03.0 (Posit team, 2023) using the *ggplot2* package (Wickham, H., 2016). Welch’s t-tests were used to compare the mean of two independent groups: cell areas and cell volumes in different cell layers, and the apparent Young’s modulus in WT and mutant SAM. When we use boxplots, the boxes extend from the first to the third quartile and the whiskers from 10% to 90% of the values, the solid black line represents the median of the distribution. Throughout the manuscript, average values are means ± s.d. On scatter plots, the regression line represents the best linear fit of the data, along with its 95% confidence interval. Pearson’s correlation coefficients and the related p-values were calculated with *ggplot2*.

### Analytical models

We used a morphoelastic framework, which involves a multiplicative decomposition of the deformation gradient into growth tensor, describing changes in mass or density, and an elastic tensor, accounting for the elastic material response, commonly used in the literature (Goriely and Ben Amar, 2005; Moulton et al., 2013). The outer L1 layer of cells in the SAM is modelled as a planar elastic rod attached to an elastic foundation that represents the inner layers (Almet et al., 2019; Moulton et al., 2013) (Fig. S2). The elastic rod grows axially in the x-direction, which generates compressive stresses due to clamped boundary conditions at the two rod ends. These stresses lead to instability, and buckling that result in the formation of folded structures, which closely resemble the shapes obtained in our experimental data. We used the model to explore differences in the growth rates and stiffness in the L1 and L2 layers of the various groups in our study (see supplementary methods for details).

### Atomic force microscopy (AFM) and data analysis

Meristems were dissected to remove all flower buds older than stage 3 the day before AFM observations and placed in ACM in 60 mm plastic Petri dishes. Acquisitions were carried out in distilled water at room temperature on a stand-alone JPK Nanowizard III microscope equipped with a CellHesion module (100 μm-range Z-piezo) using the Quantitative Imaging mode (QI). We used a silica spherical tip with a radius of 400 nm (Special Development SD-sphere-NCH, Nanosensors) mounted on a silicon cantilever with a nominal force constant of 42 N.m^-1^. Scan size was generally a square field of 45 x 45 μm with pixel size of approximately 500 nm. An applied force trigger of 1 μN was used in order to indent only the cell wall (Milani et al., 2011; Tvergaard and Needleman, 2018), with a ramp size of 2 μm, an approach speed of 10 μm.s^-1^ and a retraction speed of 100 μm.s^-1^. Apparent Young’s modulus was obtained by fitting the first 100-150 nm of the force displacement curve with a Hertz model and subsequently corrected for the effect of slope (see supplementary methods).

### RNA in situ hybridisations

RNA *in situ* hybridisations on sections were performed according to published protocols (Long et al., 1996). Meristems were dissected just after bolting, fixed in FAA (formaldehyde 3.7% [v/v], ethanol 50% [v/v], acetic acid 5% [v/v], H_2_O to final volume), washed, dehydrated and embedded in paraplast. Embedded samples were cut (10 μm-thick) and attached to pre-coated glass slides (Superfrost Plus Gold, Fisher Scientific). Antisense probes were made from PCR products using cDNA from inflorescences as a template, except for *CLV3*, which was amplified using genomic DNA (see supplementary methods for detailed primer list). Those PCR products were transcribed into RNA and then labelled using digoxigenin (DIG)-UTP. All probes were filtered on columns (CHROMA SPIN-30 columns, Clontech) to remove remaining nucleotides. Immunodetection was performed using an anti-DIG antibody coupled to alkaline phosphatase (Anti-Digoxygenin-AP, Fab fragments, Roche), whose activity was detected by the chromogenic method using NBT/BCIP (Roche). Sections were finally washed with water and observed under a Zeiss Imager M2 microscope equipped with an AxioCam Mrc camera, and 10 X or 20 X objectives in DIC (differential interference contrast) mode.

### Hormone treatments

Auxin treatments were performed by immersing dissected SAM in 1 or 2 mM Indole-3-acetic acid (IAA) solution (Sigma) for five hours as described (Galvan-Ampudia et al., 2020). *DR5* reporter expression was monitored in the samples by confocal microscopy approximately 24 hours post-treatment. Treatments were carried out on *clv1-8* (in the Col-0 accession, as in Fig. S9D, E), *clv3-2* plants (in the L*er* accession, as in Fig. S9A, B), or in F1 plants of a cross between *clv3-2* and *clv3-21* (mixed Col-0 and L*er* background, as in Fig. 6C-D, I-L).

## Acknowledgements

We thank V. Mirabet and N. Dubrulle for help with preliminary AFM measurements, A. Kiss for help with 3D segmentations and surface curvature detection, Zofia Haftek-Terreau and G. Gilbert for help with *in situ* hybridisations, and Aurélie Chauvet for help with molecular biology. We thank the PLATIM/LyMIC imaging facility of the SFR Biosciences (UMS3444/CNRS, US8/Inserm, ENS de Lyon, UCBL1). We thank A. Lacroix, J. Berger, P. Bolland, H. Leyral, and I. Desbouchages for assistance with plant growth and logistics. This article is based on LRL’s PhD thesis: Rambaud-Lavigne, L. E. S. (2018), The genetics and mechanics of stem cells at the Arabidopsis shoot apex. *PhD thesis*, University of Lyon, Lyon, France, downloadable at: theses.hal.science/tel-02366972.

## Author contributions

Conceptualization: LRL, AB, PD; Methodology: AC, NG, SB, AB; Software & Modeling: AC, NG; Validation: LRL, VB, PD; Formal Analysis: QL, AC, LRL, PD; Investigation: LRL, SB, VB, PD; Data Curation: LRL, PD; Writing - Original Draft Preparation: LRL, AC, NG, PD; Writing - Review & Editing: LRL, AC, SB, NG, AB, PD; Visualisation: LRL, AC, QL, NG, PD; Supervision: NG, AB, PD; Funding Acquisition: LRL, NG, PD.

## Data availability

All datasets and scripts will be made available before publication (online repository).

