## Supplementary data for "Heterogeneous identity, stiffness and growth characterise the shoot apex of *Arabidopsis* stem cell mutants"

### Supplementary information

Figure S1: Phenotypes of fasciated *clv1* and *clv3* SAM.

Figure S2: Schematic of the morphoelastic rod model.

Figure S3: Supplementary simulations of the analytical model.

Figure S4: Supplementary analyses of growth in WT and *clv* SAM.

Figure S5: Supplementary analyses of AFM experiments.

Figure S6: Expression of *APUM10* and *WUS* on a *clv3-2* SAM.

Figure S7: Expression of *APUM10* and *MP* on a *clv3-2* SAM.

Figure S8: Expression of *APUM10*, *WUS*, and *MP* in various *clv* alleles.

Figures S9: Effect of auxin treatment on *clv1* and *clv3* SAM.

Table S1. Mean and standard deviation of apparent Young's modulus by genotype.

Table S2. Mean and standard deviation of apparent Young's modulus organised by genotype and by scan.

Supplementary materials and methods

Supplementary references

### Figure S1

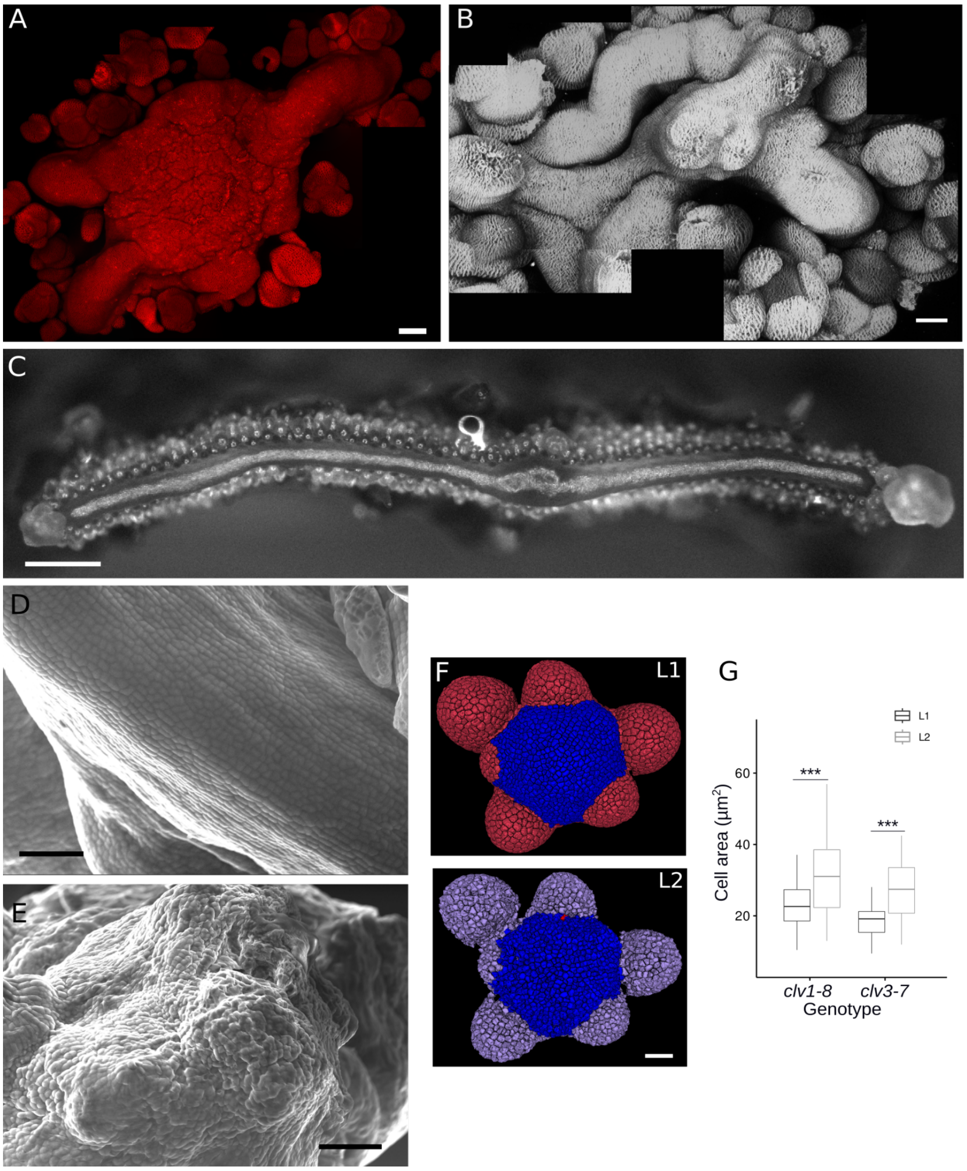


**Figure S1: Phenotypes of fasciated *clv1* and *clv3* SAM.**

**(A, B)** Maximum intensity projections of confocal stacks showing a fasciated *clv3-21* SAM crossed to the *pUBQ10::LTI6b-tdTomato* cell membrane marker (A), and a *clv3-2* SAM stained with FM4-64. Images were produced from a collage of smaller projections. **(C)** Bright field photograph of a dissected *clv1-8* SAM. **(D, E)** SEM images of the linear part (D), and an outgrowth (E) in a *clv3-2* SAM. **(F)** 3D reconstruction of a WT SAM showing L1 (top) and L2 (bottom) cells. Cells used for volume computation are in blue. **(G)** Boxplots of cell area in L1 (black) and L2 (grey) cells of *clv1-8* and *clv3-7* SAM. Mean ± s. d. areas are 23.0 ± 5.8 µm$^2$ (*clv1-8* L1) and 31.0 ± 10.3 µm$^2$ (*clv1-8* L2), 18.7 ± 4.1 µm$^2$ (*clv3-7* L1) and 26.8 ± 7.9 µm$^2$ (*clv3-7* L2). L1 and L2 cells have significantly different areas in both mutants (Welch's t-test, p = 1.35e-7 (*clv1-8*); p = 2.98e-6 (*clv3-7*)). n = 133 (L1) and 64 (L2) cells from 1 *clv1-8* SAM, 40 (L1) and 33 (L2) cells from 1 *clv3-7* SAM. Scale bars: 50 µm (A, B), 1 mm (C), 25 µm (D, E), 20 µm (F).

### Figure S2

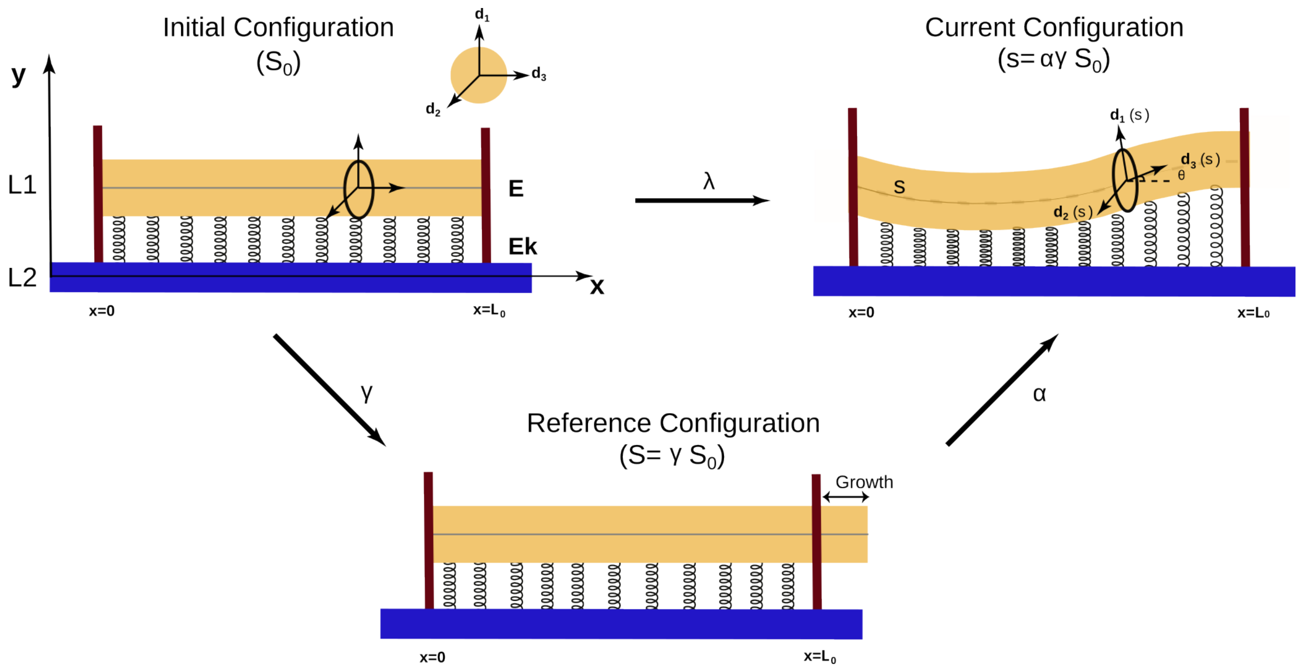


**Figure S2: Schematic of the morphoelastic rod model.**

Schematic of adopted morphoelastic framework of a planar growing rod attached to an elastic foundation. The initial configuration shows the rod attached to the foundation before it starts to grow. The reference configuration is the unstressed virtual configuration where the rod is allowed to grow before being subjected to boundary conditions. The current configuration is where the growing rod on the foundation is subjected to the boundary conditions and the stresses, which induces deformation. The growth, elastic and total stretches are denoted by γ, α and λ respectively. The growing rod is embedded in the x - y plane, with the local orthonormal basis denoted as ($d_1$, $d_2$, $d_3$).

### Figure S3

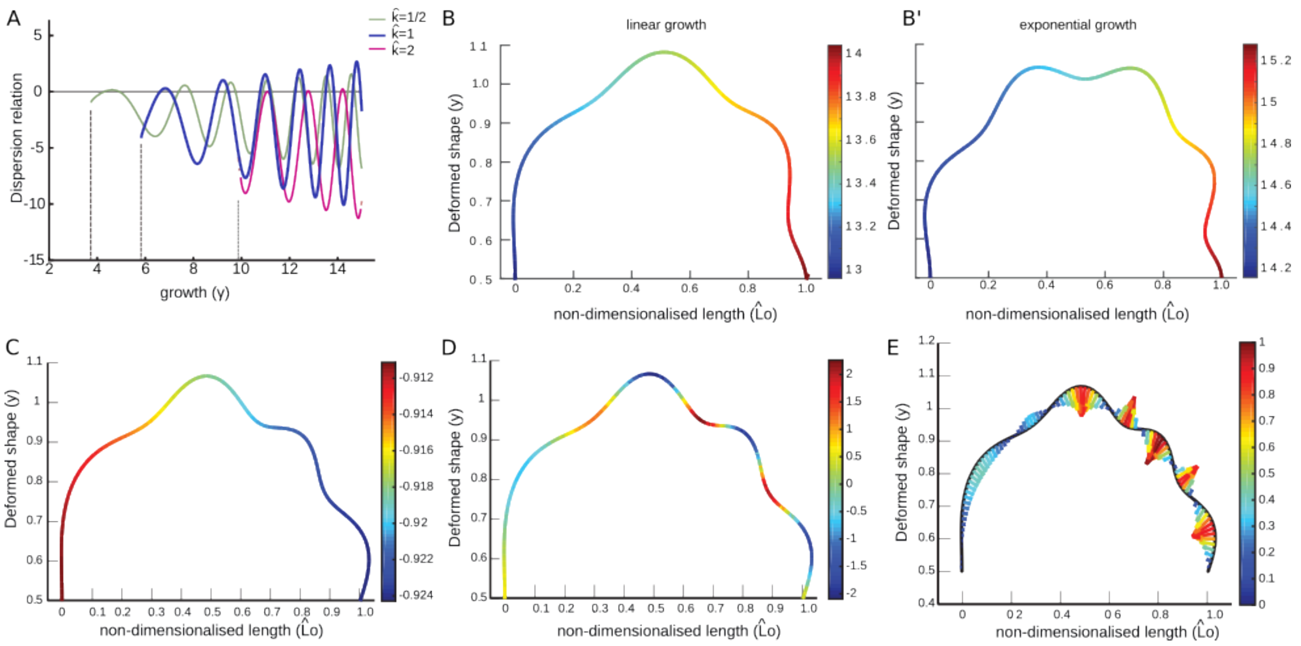


**Figure S3: Supplementary simulations of the analytical model.**

**(A)** Effect of underlying foundation stiffness ($\hat{k}$) on critical growth rates (γ*), which alters the buckled shapes. **(B, B')** Deformed shape obtained as a result of growth-induced buckling in the case of linear (left) and exponential (right) growth laws. **(C, D)** Growth-induced buckling causes changes in mechanical responses – growth-induced compressive stresses (C) and bending moment (D). Spatial distribution of these responses are plotted for a cosine form of growth law. **(E)** The changes in curvature of the SAM shapes corroborate with the spatial distribution of bending moments across the SAM, which depend on growth rates.

### Figure S4

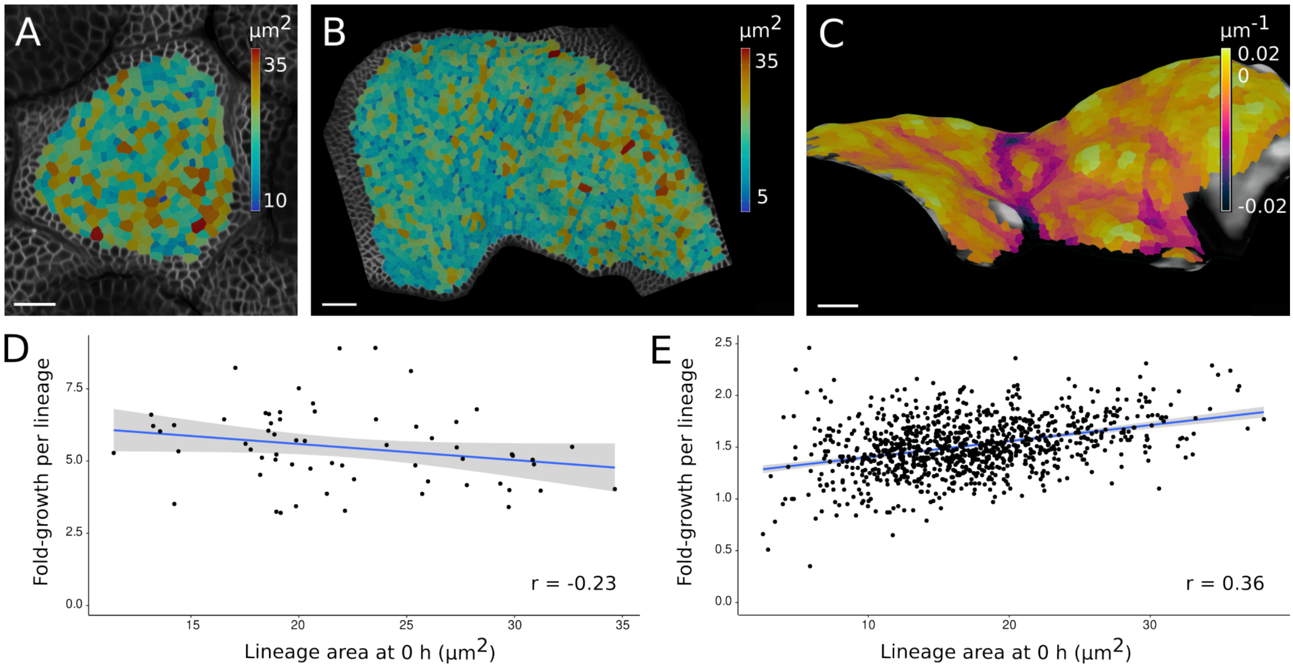


**Figure S4: Supplementary analyses of growth in WT and *clv* SAM.**

**(A, B)** Heat maps of initial cell lineage area in the WT Col-0 (A) and the *clv3-7* (B) SAM shown in Fig. 3. **(C)** Side view of the curvature map shown in Fig. 3H. **(D, E)** Scatter plots of initial lineage area as a function of the fold-growth per lineage in WT (D) and *clv3-7* (E) SAM. Regression lines representing the best fit are shown in blue, and confidence intervals as shaded regions around them. (r) Pearson's correlation coefficients. *P*-values: $p = 0.075$ (D); $p < 0.001$ (E). Colourmap: Turbo (A, B); warm helix (C). Scale bars: 20 μm (A-C).

### Figure S5

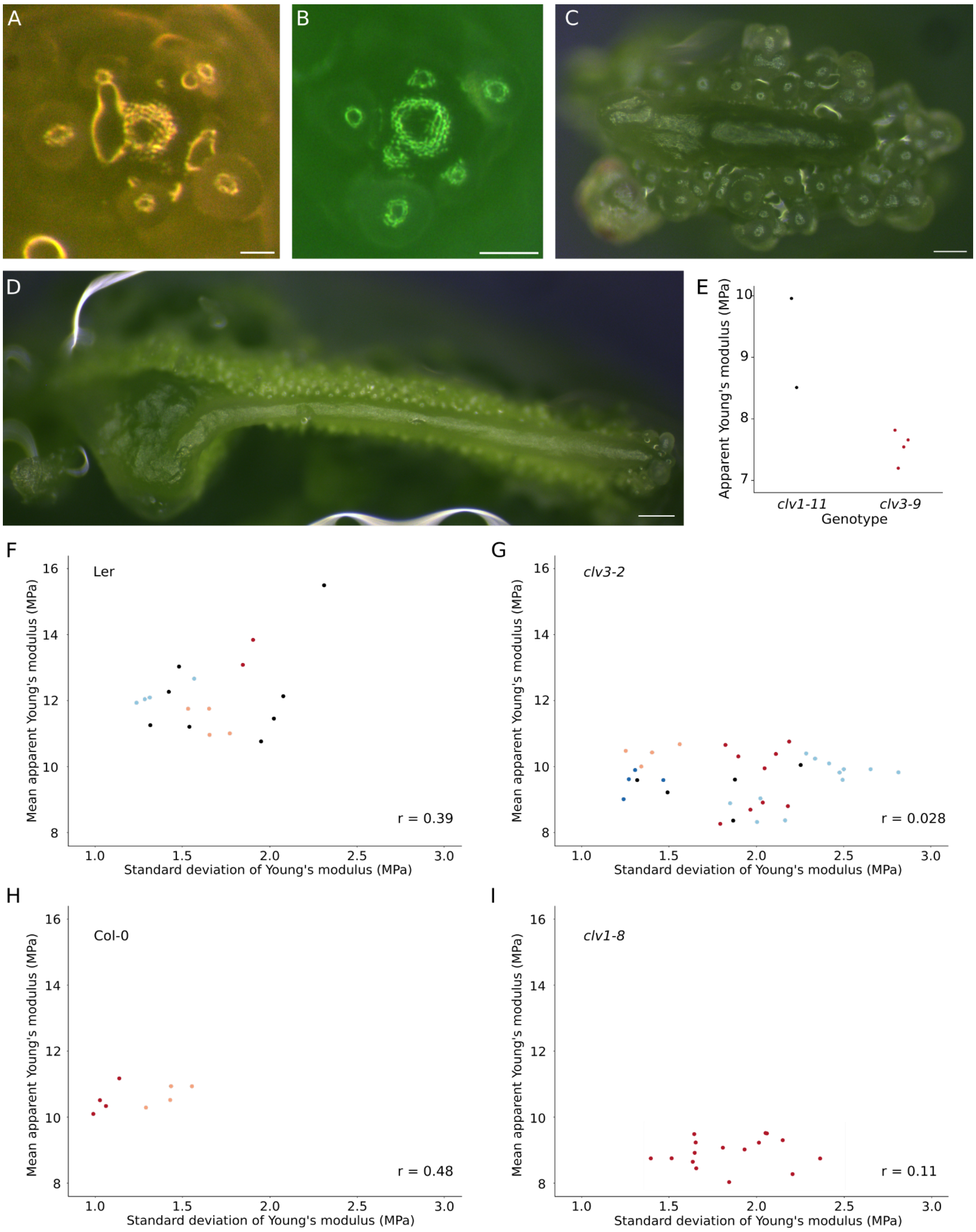


**Figure S5: Supplementary analyses of AFM experiments.**

**(A-D)** Bright field images of representative L*er* (A), Col-0 (B), *clv3-2* (C), and *clv1-8* (D) SAM used for AFM experiments. **(E)** Mean apparent Young's moduli measured on epidermal cells of *clv1-11*, and *clv3-9* SAM. Compare *clv1-11* to L*er*, and *clv3-9* to Col-0 (Fig. 3G). **(F-I)** Scatter plots of the mean apparent Young's modulus per scan as a function of its standard deviation in L*er* (F), *clv3-2* (G), Col-0 (H), and *clv1-8* (I). These panels use the same data and colour-code as in Fig. 4G, H. (r) Pearson's correlation coefficients. *P*-values: $p = 0.11$ (F); $p = 0.88$ (G), $p = 0.23$ (F), $p = 0.68$ (F). n = 2 scans on 2 samples (*clv1-11*), n = 4 non-overlapping scans on 1 SAM (*clv3-9*). Scale bars: 500 µm (A-D).

#### Figure S6

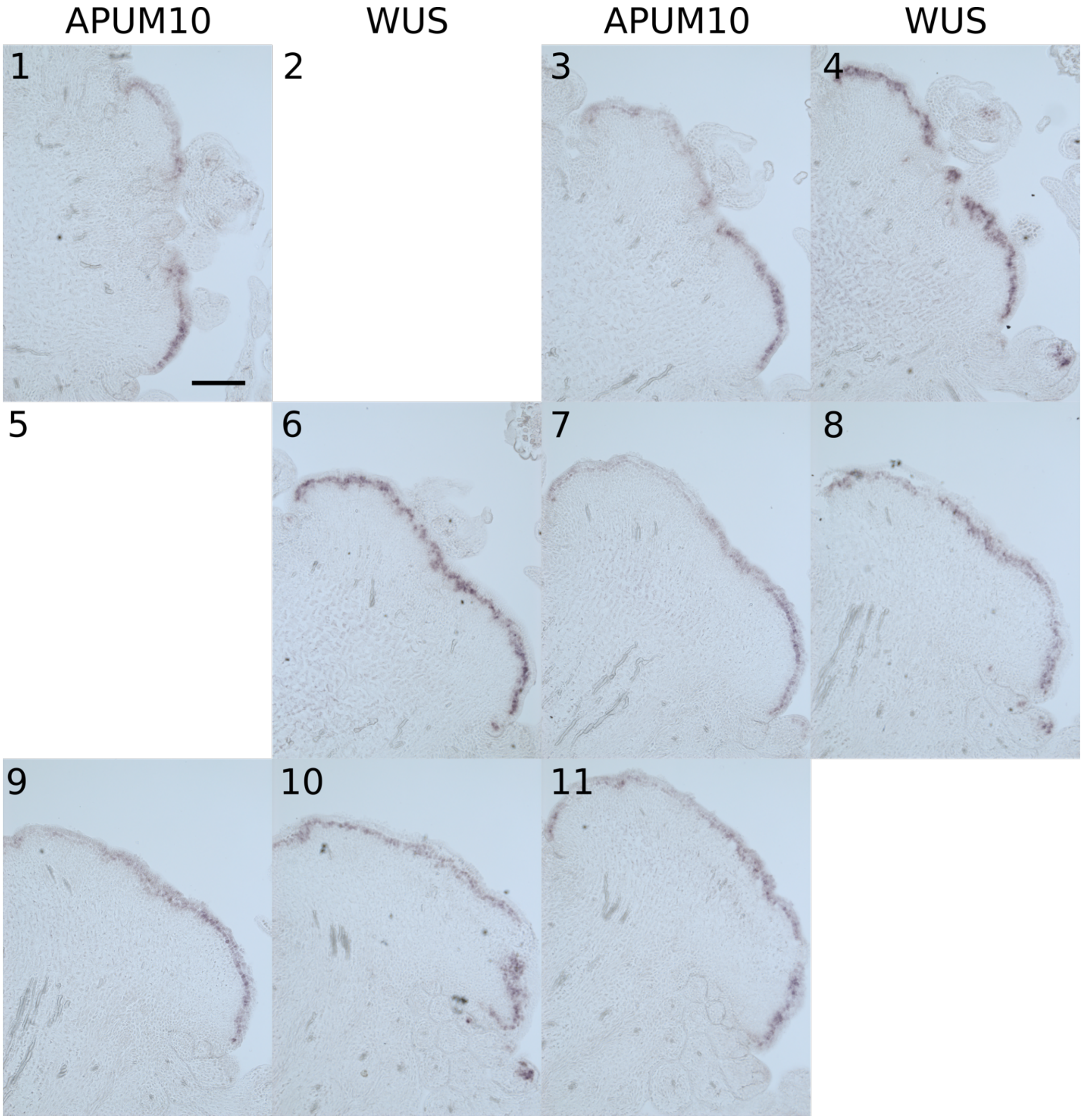


**Figure S6: Expression of *APUM10* and *WUS* on a *clv3-2* SAM.**

Consecutive sections of a single *clv3-2* SAM hybridised with *APUM10* (odd numbers) and *WUS* (even numbers) probes. Section thickness: 10 µm. Sections 2 and 5 are missing in the series due to damage. Scale bar: 100 µm (shown on section 1).

**Figure S7**

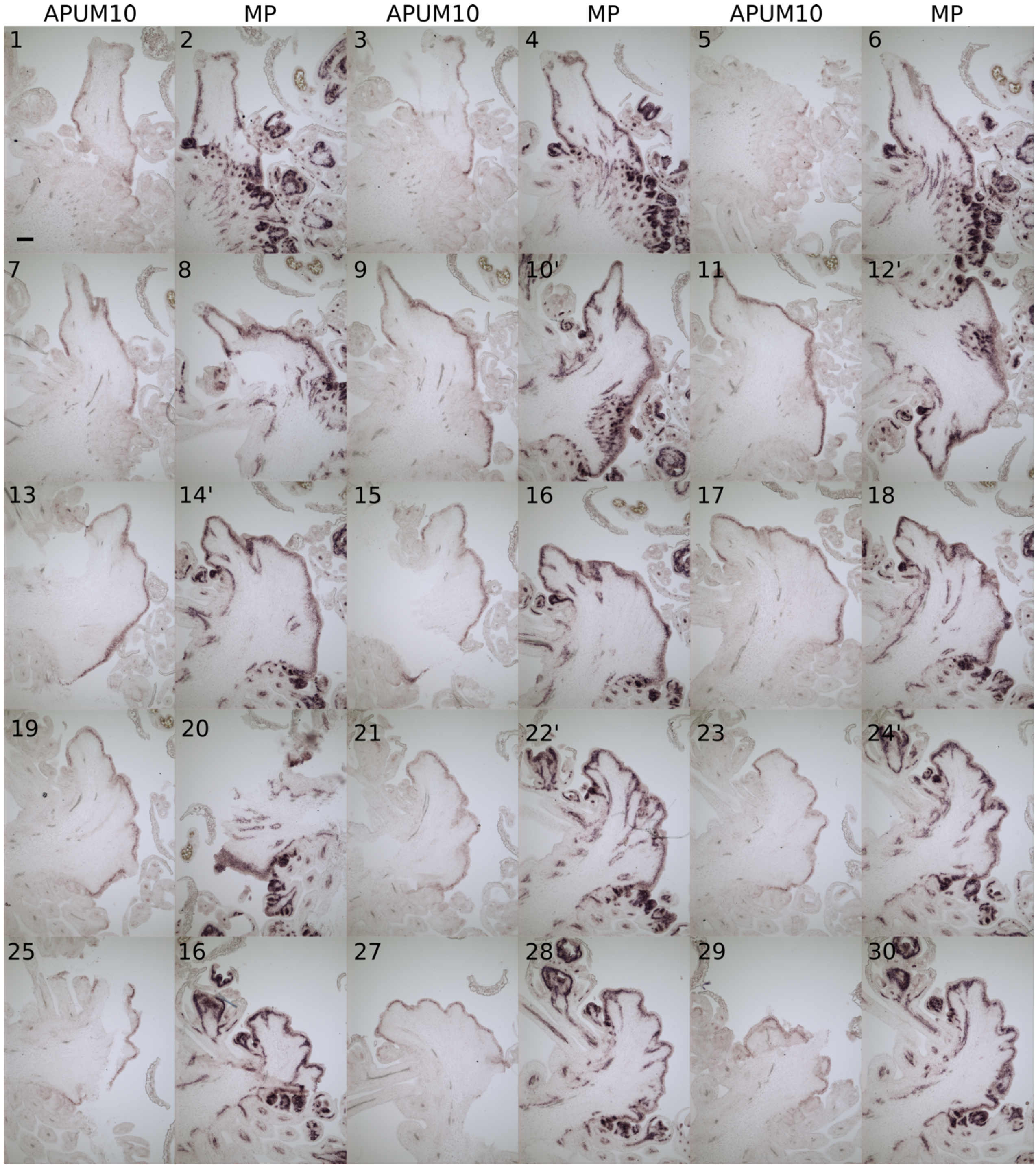


**Figure S7: Expression of *APUM10* and *MP* on a *clv3-2* SAM.**

Consecutive sections of a single *clv3-2* SAM hybridised with *APUM10* (odd numbers) and *MP* (even numbers) probes. Section thickness: 10 µm. Scale bar: 100 µm (shown on section 1).

#### Figure S8

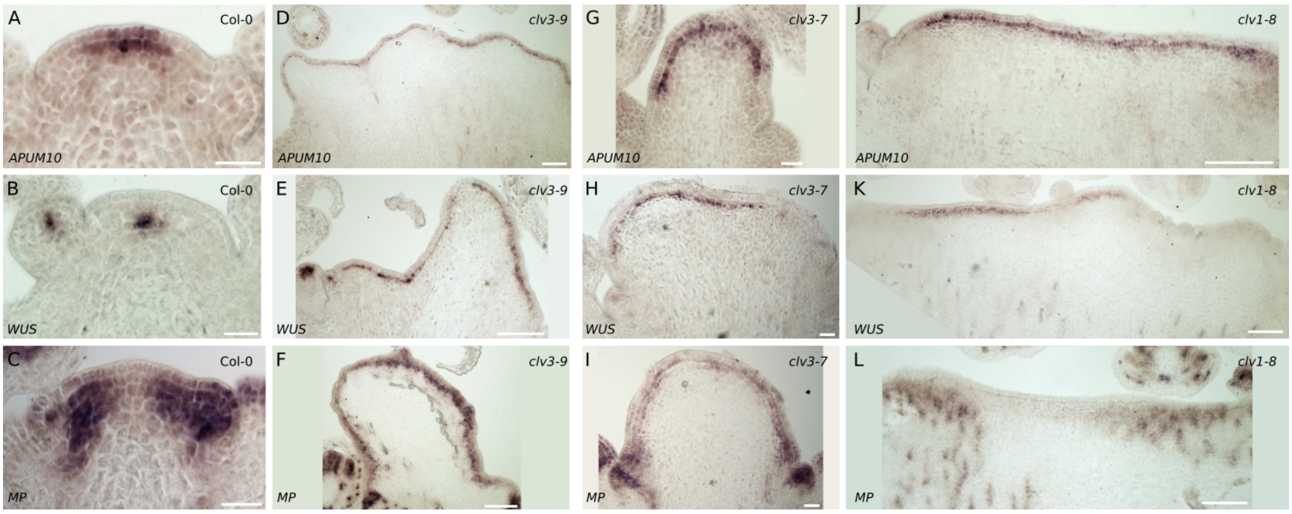


**Figure S8: Expression of *APUM10*, *WUS*, and *MP* in various *clv* alleles.**

RNA *in situ* hybridisations showing *APUM10* (A, D, G, J), *WUS* (B, E, H, K), and *MP* (C, F, I, L) probes hybridised on Col-0 (A-C), *clv3-9* (D-F), *clv3-7* (G-I), and *clv1-8* (J-L) SAM. Section thickness: 10 µm. Scale bars: 25 µm (A-C, G-I), 100 µm (D-F, J-L).

### Figure S9

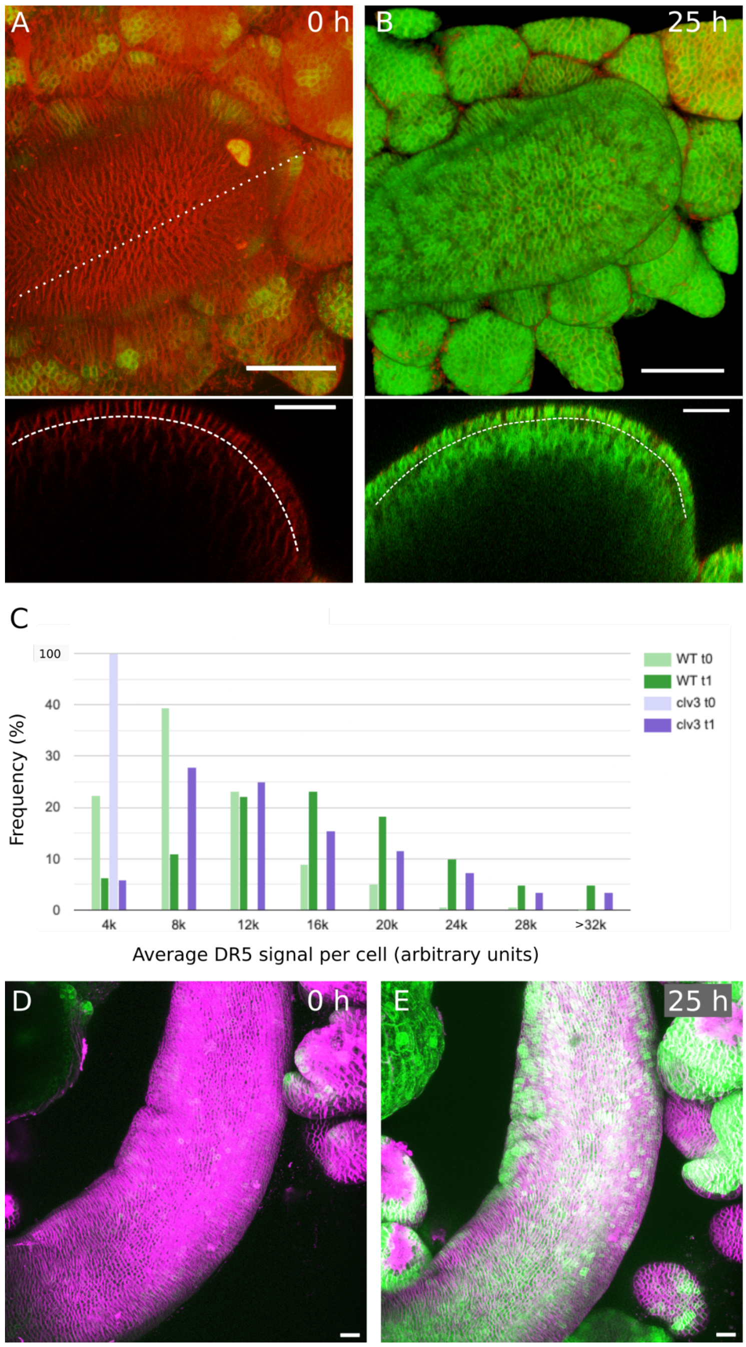


**Figures S9: Effect of auxin treatment on *clv1* and *clv3* SAM.**

**(A, B)** Projections of confocal stacks acquired before (A) and after (B) IAA treatment of a *clv3-2* SAM expressing the *DR5::GFPer* reporter (n=1/11). Plasma membranes are dyed with FM4-64. An orthogonal view of the SAM is taken along the dotted line shown in (A). Dashed lines in orthogonal sections represent the interface between L1 and L2 layers. **(C)** Histogram of the average DR5 signal per cell in WT and *clv3* meristems (samples shown in Fig. 6E-L) before (t0) and after (t1) auxin treatment. **(D, E)** Projections of confocal stacks acquired before (D) and after (E) IAA treatment of a *clv1-8* SAM expressing the *pUBQ10::Lti6b-tdTomato* and *DR5::GFPer* reporters (n=5/5). Scale bars: 50 µm (top views in A, B), 30 µm (orthogonal sections in A, B), 20 µm (D, E).

**Table S1.** Mean and standard deviation of apparent Young's modulus by genotype.

| Genotype | Y. m. (MPa) | s. d. (MPa) |
|---|---|---|
| L*er* | 12.16 | 1.17 |
| *clv3-2* | 9.63 | 0.74 |
| *clv1-11* | 9.21 | 1.04 |
| Col-0 | 10.58 | 0.38 |
| *clv1-8* | 8.94 | 0.45 |
| *clv3-9* | 7.56 | 0.28 |

Y. m.: mean apparent Young's modulus

s. d.: standard deviation

**Table S2.** Mean and standard deviation of apparent Young's modulus organised by genotype and by scan.

| L*er* | | | *clv3-2* | | | Col-0 | | | *clv1-8* | | |
|---|---|---|---|---|---|---|---|---|---|---|---|
| scan | Y. m. (MPa) | s. d. (MPa) | scan | Y. m. (MPa) | s. d. (MPa) | scan | Y. m. (MPa) | s. d. (MPa) | scan | Y. m. (MPa) | s. d. (MPa) |
| 10.00.36.578 | 12.27 | 1.42 | 14.37.52.713 | 9.20 | 1.49 | 11.36.13.473 | 10.08 | 0.99 | 13.41.24.492 | 8.75 | 2.36 |
| 11.33.38.262 | 11.46 | 2.02 | 15.52.09.686 | 9.60 | 1.88 | 11.40.12.309 | 10.49 | 1.03 | 13.46.07.197 | 8.04 | 1.84 |
| 12.46.23.885 | 15.50 | 2.31 | 10.05.19.069 | 8.36 | 1.87 | 11.44.30.666 | 10.32 | 1.06 | 13.50.10.421 | 8.28 | 2.20 |
| 10.35.34.897 | 10.77 | 1.95 | 14.07.57.548 | 9.58 | 1.32 | 11.48.08.104 | 11.15 | 1.14 | 14.02.56.630 | 9.51 | 2.06 |
| 10.12.00.678 | 11.26 | 1.31 | 13.51.25.419 | 10.04 | 2.26 | 17.20.46.317 | 10.27 | 1.29 | 14.13.20.405 | 8.66 | 1.63 |
| 13.30.45.041 | 13.04 | 1.48 | 14.24.13.297 | 10.38 | 2.12 | 17.24.25.083 | 10.50 | 1.43 | 14.21.39.713 | 8.93 | 1.65 |
| 12.05.49.800 | 11.21 | 1.54 | 14.59.23.378 | 8.91 | 2.04 | 17.28.06.472 | 10.92 | 1.55 | 14.25.35.938 | 9.08 | 1.81 |
| 10.52.07.813 | 12.14 | 2.08 | 15.12.59.108 | 8.80 | 2.18 | 17.31.45.141 | 10.92 | 1.43 | 14.29.38.142 | 9.24 | 1.65 |
| 11.48.01.525 | 13.09 | 1.84 | 15.16.38.202 | 8.26 | 1.80 | | | | 14.41.13.282 | 9.31 | 2.15 |
| 13.45.39.389 | 13.84 | 1.90 | 15.20.34.843 | 8.69 | 1.97 | | | | 14.45.26.192 | 9.03 | 1.93 |
| 13.08.50.178 | 10.97 | 1.65 | 15.28.55.701 | 9.94 | 2.05 | | | | 14.51.28.489 | 9.53 | 2.05 |
| 13.12.28.125 | 11.01 | 1.77 | 15.44.29.657 | 10.75 | 2.19 | | | | 14.55.07.380 | 9.24 | 2.01 |
| 13.16.42.713 | 11.76 | 1.65 | 15.48.15.971 | 10.65 | 1.83 | | | | 15.02.14.295 | 8.76 | 1.51 |
| 13.21.13.360 | 11.76 | 1.53 | 15.53.44.955 | 10.30 | 1.90 | | | | 15.06.10.958 | 8.46 | 1.65 |
| 17.01.52.144 | 12.05 | 1.28 | 15.53.09.955 | 10.47 | 1.25 | | | | 15.09.56.555 | 8.76 | 1.40 |
| 17.05.35.944 | 12.10 | 1.31 | 15.57.06.383 | 10.67 | 1.56 | | | | 15.13.43.543 | 9.49 | 1.64 |
| 17.09.15.786 | 11.94 | 1.23 | 16.00.42.679 | 10.42 | 1.41 | | | | | | |
| 17.13.15.602 | 12.67 | 1.57 | 16.04.22.968 | 10.00 | 1.34 | | | | | | |
| | | | 16.11.30.019 | 8.32 | 2.01 | | | | | | |
| | | | 16.15.10.858 | 8.88 | 1.85 | | | | | | |
| | | | 16.19.27.862 | 9.03 | 2.03 | | | | | | |
| | | | 16.23.07.691 | 8.37 | 2.17 | | | | | | |
| | | | 16.33.42.658 | 10.39 | 2.29 | | | | | | |
| | | | 16.37.25.298 | 9.91 | 2.66 | | | | | | |
| | | | 16.41.06.047 | 9.81 | 2.48 | | | | | | |
| | | | 16.44.48.434 | 9.59 | 2.50 | | | | | | |
| | | | 16.52.53.170 | 9.92 | 2.50 | | | | | | |
| | | | 16.56.32.407 | 10.24 | 2.34 | | | | | | |
| | | | 17.00.14.200 | 10.09 | 2.42 | | | | | | |
| | | | 17.03.51.615 | 9.82 | 2.82 | | | | | | |

| Ler | | | clv3-2 | | | Col-0 | | | clv1-8 | | |
|---|---|---|---|---|---|---|---|---|---|---|---|
| scan | Y. m. (MPa) | s. d. (MPa) | scan | Y. m. (MPa) | s. d. (MPa) | scan | Y. m. (MPa) | s. d. (MPa) | scan | Y. m. (MPa) | s. d. (MPa) |
| | | | 18.00.36.632 | 9.59 | 1.47 | | | | | | |
| | | | 18.04.14.726 | 9.01 | 1.24 | | | | | | |
| | | | 18.07.59.800 | 9.89 | 1.31 | | | | | | |
| | | | 18.11.45.950 | 9.61 | 1.27 | | | | | | |

Y. m.: mean apparent Young's modulus

s. d.: standard deviation

#### Supplementary materials and methods

##### Plant material

The following plant lines were used in supplementary data: *clv3-9* (Nimchuk et al., 2015), *clv1-11* (Dievart, 2003).

##### Primers used to generate RNA *in situ* hybridisation probes

*At1G35750* (*APUM10*): BP840 5'-TTC CGA TCA TCG TCG TCT TC-3', and BP841 5'-TGT AAT ACG ACT CAC TAT AGG GCT GCA ATA AGG ATT CGA CTG TAG-3'

*At2G27250* (*CLV3*): BP637 5'-ATG TCC GGT CCA GTT CAA CAA C-3', and BP638 5'-TGT AAT ACG ACT CAC TAT AGG GCG GTC AGG TCC CGA AGG AAC A-3'

*At1G19850* (*MP*): BP1059 5'-CAC CAT GAT GGC TTC ATT GTC T-3', and BP1060 5'-TAA TAC GAC TCA CTA TAG GGT GAA ACA GAA GTC TTA AGA TCG

*At2G17950* (*WUS*): BP663 5'-CAA CAA GTC CGG CTC TGG TG-3', and BP664 5'-TGT AAT ACG ACT CAC TAT AGG GCG GGA AGA GAG GAA GCG TAC GTC G-3'

##### Analytical models

*Morphoelastic framework of growing rod on elastic foundation*

We delineate three distinct configurations: first, an 'initial' configuration parametrised by the arc length, $S_0$. Second, a grown virtual configuration, referred to as the 'reference' configuration, parametrised in terms of *S*, and finally, the 'current' configuration of the deformed rod under applied stress, parameterised using $s$ (Fig. S2). The rod with an initial length, $L_0$, is constrained between two supports, and grows such that the growth stretch is given by $\gamma = \frac{\partial S}{\partial S_0}$, and the elastic stretch by $\alpha = \frac{\partial s}{\partial S}$ . The total stretch, λ, in the rod from the initial to current configurations is given by

$\lambda = \alpha\gamma = \frac{\partial s}{\partial S_0} = \frac{\partial s}{\partial S}\frac{\partial S}{\partial S_0}$, (1).

*Geometry and Mechanics of the growing rod*

We adapt a theoretical framework developed earlier (Almet et al., 2019) to define the geometric and mechanical descriptors for the rod on the elastic foundation. We use Frenet-Serret equations (Coleman et al., 1993) to define the rod geometry, which is only a function of the material parameter - arc length *S*. We simplify the equations by assuming the rod to be unshearable. Using equilibrium equations (for linear and angular momentum balance), we determine the constitutive relations for the growing rod on an elastic foundation in x-y plane (Almet et al., 2019; Goriely and Ben Amar, 2005; Moulton et al., 2013). The bending moment (m) is related to curvature by

$m = \frac{EI}{\gamma}\frac{\partial\theta}{\partial S_0}$, (2).

where E represents the elastic modulus of the rod, I is the moment of inertia, and θ is the angle of the tangent basis vector with the horizontal axis. The axial stresses are linked to the elastic stretch, α, in a linear elastic framework by

$F cos\theta + G sin\theta = EA(\alpha - 1)$, (3).

where F and G are the resultant forces along the x and y axis, and A is the area of cross-section of the elastic rod. The body force acting on the rod due to the underlying foundation is calculated by

$\gamma f = \gamma(f\, e_x + g\, e_y) = -Ek[(x(S_0) - S_0)\, e_x + (y(S_0) - y_0) e_y]$, (4).

where *k* is a non-dimensional parameter that represents the ratio of stiffness of the foundation to that of the rod. The modulus of the foundation is given as '*Ek*' and the rod is assumed to be clamped at both ends and to have an initial length $L_0$. The boundary conditions are hence given as

$x(0) = 0, x(L_0) = 0, y(0) = 0, y(L_0) = 0, \theta(0) = 0, \theta(L_0) = 0$, (5).

*Linear stability analysis to obtain critical growth stretch ($\gamma$*) for the morphoelastic rod*

We non-dimensionalise the governing equations using standard Kirchhoff scaling (Almet et al., 2019; Goriely and Tabor, 1997) and obtain the following scaling relationships:

$\{S_0, x, y\} = \left(\frac{A}{I}\right)^{\frac{1}{2}}\{\hat{S}_0, \hat{x}, \hat{y}\}, \{F, G\} = EA\{\hat{F}, \hat{G}\}, m = E(AI)^{\frac{1}{2}}\{\hat{m}\}$, (6).

The constitutive equation is hence given by

$F cos\theta + G sin\theta = (\alpha - 1)$, (7a).

$\gamma m = \frac{\partial\theta}{\partial S_0}$, (7b).

and the non-dimensionalised clamped boundary conditions are given by

$x(0) = 0, x(\widehat{L_0}) = 0, y(0) = 0, y(\widehat{L_0}) = 0, \theta(0) = 0, \theta(\widehat{L_0}) = 0$, (8).

where the dimensionless parameters are

$\widehat{L_0} = \left(\frac{A}{I}\right)^{\frac{1}{2}} L_0$ and $\hat{k} = k\frac{I}{A^2}$, (9).

We perform a linear stability analysis to obtain the critical growth stretches, $\gamma^*$, and the corresponding buckling modes obtained using the model.

We expand the terms such that $x = x_0 + \delta x_1 + O(\delta^2)$ for an arbitrary small parameter, $\delta$, and consider O(δ) terms alone for the linearized system. The deformed shape of the curved rod is given by

$y^1 = C_1[cos(\omega_+ S_0) - cos(\omega_- S_0) + C_2 sin(\omega_+ S_0) + C_3 sin(\omega_- S_0)]$, (10).

where $C_2 = \frac{\omega_-(\cos(\omega_-\widehat{L_0}) - \cos(\omega_+\widehat{L_0}))}{\omega_- \sin(\omega_+\widehat{L_0}) - \omega_+ \sin(\omega_-\widehat{L_0})}$, $C_3 = \frac{\omega_+(\cos(\omega_+\widehat{L_0}) - \cos(\omega_-\widehat{L_0}))}{\omega_- \sin(\omega_+\widehat{L_0}) - \omega_+ \sin(\omega_-\widehat{L_0})}$, (11a).

$\omega_\pm = \alpha \pm (a^2 - b^2)$, (11b)

$\alpha = \frac{\gamma - 1}{2}$ and $b = \sqrt{\hat{k}y}$, (11c).

where $C_1$ is the amplitude, $\gamma$ is the growth of the rod and $k$ is the foundation stiffness. The initial rod is curved and its ordinates are denoted by $y^0$. The final deformed shape is obtained by the superposition of $y^0$ and $y^1$.

The critical growth ($\gamma^*$) satisfies the relation:

$a\, sin(\omega_+\widehat{L_0})\, sin(\omega_-\widehat{L_0}) + b\, cos(\omega_+\widehat{L_0})\, cos(\omega_-\widehat{L_0}) - b = 0$, (12).

Solutions to this equation provide values of the various buckling modes that are characterised by the critical growth, $\gamma^*$, for various stiffness values considered in this study (Fig. S3A). The critical $\gamma^*$ values are next used to obtain the various buckled shapes. The smallest real value of $\gamma$>1, $\gamma^*_{inf}$, that satisfies the above relation occurs when a=b.

$\gamma^*_{inf} = 1 + 2\hat{k} + 2(\hat{k} + \hat{k}^2)^{\frac{1}{2}}$, (13).

*Asymmetric buckled shapes and the role of growth and stiffness heterogeneities*

To examine the effects of local growth heterogeneities on buckling, we differentially distribute growth along the planar rod. To this end, we use three different distributions of growth over the length of the rod – characterised by cosine, linear and exponential equations given by (Fig. 2C)

$\gamma = \gamma_0\left(1 - \varepsilon cos\left(\pi\frac{S_0}{\widehat{L_0}}\right)\right)$ Cosine

$\gamma = \gamma_0\left(1 + \varepsilon\left(S_0 - \frac{\widehat{L_0}}{2}\right)\right)$ Linear, (14).

$\gamma = \gamma_0\left(1 + \varepsilon e^{\left(S_0 - \frac{\widehat{L_0}}{2}\right)}\right)$ Exponential

where $\varepsilon$ is a very small number and $\gamma_0$ is set to a number above the critical value (e.g. 12.23). Fig. 2D and Fig. S3 B-B' show that the number of buckling modes differ based on the choice of growth law irrespective of the same initial value of $\gamma_0$ ($\hat{k}$=1 for all cases). We compare the shapes and buckling modes obtained from the models with experiments on longitudinal sections in various *clv* mutant SAM. We also investigate the effect of variable stiffness on differential buckling in our model. To do this, we replace the constant stiffness ratio ($\hat{k}$) between the rod and foundation, with a spatial distribution of stiffness ($\hat{k} = \widehat{k_0} f(S_0)$) over the length of the meristem, and perform the analyses for a constant value of $\gamma_0$ . Specifically, a cosine distribution is used, given by

$\hat{k} = \hat{k}_0(1 + \epsilon\, cos(\pi \frac{S_0}{\widehat{L_0}}))$, (15).

Where $\hat{k}_0$=1 and $\gamma_0$=12.23 respectively. We observe that variability in stiffness across the length can also induce asymmetric buckling (Fig. 2E).

*Variation in growth-induced compressive stresses, curvature and bending moment*

We used non-dimensionalised expressions to obtain the net compressive stresses

$F cos\theta + G sin\theta = (\alpha - 1) = \frac{1-\gamma}{\gamma}$, (16a).

And the non-dimensionalised bending moments are calculated as

$m = \frac{1}{\gamma}\frac{\partial \theta}{\partial S_0} = \frac{1}{\gamma}\frac{\partial^2 y}{\partial {S_0}^2}$, (16b).

Fig. S3C shows the distribution of compressive stresses due to growth heterogeneity along the length of the deformed rod. We compute the distribution of bending moments using the cosine form of spatial distribution of growth (Eqn .1) and have plotted them in Fig. S3D. We observe that the distribution of bending moments along the meristem length closely corroborate with the variation of curvature (Fig. S3E) with the magnitude of these quantities represented by a colormap.

**Atomic force microscopy (AFM) and data analysis**

Cantilever calibration was performed following the standard thermal noise method. We measured the deflection sensitivity by doing a linear fit of the contact part of a force curve acquired on sapphire in phosphate-buffered saline (PBS). Then, we determined the spring constant by acquiring the thermal noise spectrum of the cantilever and fitting the first normal mode peak using a single harmonic oscillator model. When possible, the same tip was used in experiments over several days. In order to reduce the offsets in force that can be introduced by each new calibration, especially by the measurements of the deflection sensitivity, we followed the SNAP protocol (Schillers et al., 2017).

Data analysis was done using JPK Data Processing software 6.0. Force vs height curves were first flattened by removing the result of a linear fit done over a portion of the baseline, in order to set this part to 0 force. A first estimation of the point of contact (POC), was obtained considering the first point crossing the 0 of forces, starting from the end of the approach curve (i.e. trigger force position). The force vs. tip-sample distance was then obtained calculating a new axis of distances as Height [m] – tip deflection Δz [m]. Young's modulus was obtained by fitting the entire force vs tip-sample distance curves (note that we used approach curves) with a Hertz model. For our analysis, we used a tip radius R of 400 nm and a Poisson's ratio $\nu$ of 0.5 (as it is conventionally set for biological material in the literature (Kulkarni et al., 2018)), where the Young's modulus, the POC and an offset in force were kept as free parameters of the fit. A preliminary filtering step was performed to exclude data points associated with $\alpha$, the angle between the normal to the surface and the z direction, superior or equal to 55°. This threshold was set based on the distribution of angles in all samples. In addition, a non systematic second filtering step was applied on samples containing artefacts on their surface, such as dust particles. The threshold criteria aims at removing data that are not well processed by the Hertz model. Thus, the data with the 10% highest residuals root mean square values were excluded.

Finally, we removed or reduced the effect of the local slope in our calculations by adapting a formula from the literature (Routier-Kierzkowska et al., 2012), thus leading to the following corrected apparent Young's modulus equation:

$E_n = E_z(1 + p^2)^{\frac{5}{4}}$, (17).

With $E_n$ the corrected apparent Young's modulus in the direction normal to the surface, $E_z$ the Young's modulus in the $z$ direction, and $p = tan(\alpha)$, with $\alpha$ being the angle between the normal to the surface and the z direction.
